# Equalizing epigenetically imprinted centromeres in early mammalian embryos

**DOI:** 10.1101/2022.10.27.514094

**Authors:** Gabriel Manske, Kelsey Jorgensen, Binbin Ma, Mansour Aboelenain, Catherine Tower, Saikat Chakraborty, Rajesh Ranjan, Arunika Das, Michael A. Lampson, Ben E. Black, Karen Schindler, Xin Chen, Saher Sue Hammoud

**Author notes:** equal contribution (Kelsey Jorgensen, Binbin Ma, Mansour Aboelenain, Catherine Tower).

## Abstract

The CENP-A histone variant epigenetically defines centromeres, where its levels and locations are precisely maintained through mitotic cell divisions. However, differences in centromere CENP-A propagation in soma versus female/male germline remains poorly understood. Here, we generated C*enpa*^*mScarlet*^ mice and followed CENP-A dynamics in gametes, zygotes, and embryos. We found that, unlike somatic cells, progenitor female and male germ cells carry high centromeric CENP-A levels that decrease upon terminal differentiation. The reduction in CENP-A is differentially regulated between sexes, resulting in a ten-fold higher level in oocytes compared to sperm. In the zygote, the parent-of-origin CENP-A asymmetry is equalized prior to initial S-phase by redistribution of nuclear CENP-A from maternal to paternal chromosomes. Redistribution of CENP-A requires both CDK1/2 and PLK1 centromeric machinery. These experiments provide direct evidence for resetting of epigenetically imprinted centromeres in early pronuclear stage embryos and imply a mechanism to sense the non-equivalency of parental chromosomes.

**Highlights:**

- Increased CENP-A density at centromeres is a conserved property of germline stem cells while CENP-A reduction is coincident with germ cell differentiation
- Paternal and maternal CENP-A containing nucleosomes are intergenerationally inherited
- CENP-A density at centromeres differs between male and female mature gametes
- Upon fertilization, maternal nuclear CENP-A is redistributed to equalize with parental CENP-A
- CENP-C and MIS18BP1 are asymmetrically enriched in the parental pronuclei in accordance with CENP-A asymmetry.
- Licensing for centromere equalization begins before zygotic DNA replication

## Introduction

Genome replication and segregation are fundamental processes that safeguard organismal development. In both mitosis and meiosis, the centromere region functions as an organizing node for spindle microtubules and the kinetochore to ensure faithful chromosome segregation. Centromeric regions are characterized by the incorporation of nucleosomes containing the histone H3 variant named centromere protein A (CENP-A), and the enrichment of the centromere satellite sequences (Palmer et al., 1991). Although centromere function is largely independent of DNA sequence, it is epigenetically defined by CENP-A nucleosomes (McKinley and Cheeseman, 2016). CENP-A-containing nucleosomes are extremely stable during the cell cycle, with a near-zero turnover rate and a precise halving dilution between centromeric sister strands during DNA replication (Bodor et al., 2013; Hemmerich et al., 2008; Jansen et al., 2007). This stability is believed to be conferred by CENP-A’s direct binding partners CENP-B, CENP-C, CENP-N, and HJURP (Fachinetti et al., 2015; Falk et al., 2015; Guo et al., 2017; Pentakota et al., 2017; Zasadzińska et al., 2018), which are in turn stabilized by interactions with the constitutive centromere-associated network (CCAN), a 16 protein heterooligomeric structure that forms the inner kinetochore (Foltz et al., 2006; Pesenti et al., 2022). After cell division, CENP-A levels are restored via pre-existing CENP-A nucleosomes, which direct new CENP-A deposition at the end of telophase and the beginning of G1 in all examined mammalian cell models (Boyarchuk et al., 2014; Jansen et al., 2007). This one-to-one mechanism of CENP-A nucleosome incorporation and maintenance ensures a tightly regulated propagation of centromeric chromatin, preserving both its genomic location and quantitative nucleosome count during mitotic cell division.

CENP-A deposition and maintenance is highly regulated across the cell cycle. It requires at least four core factors: two MIS18 subunits (MIS18α and MIS18β), the Mis18-binding protein 1 (MIS18BP1, also known as KNL2), and the Holliday Junction Recognition Protein (HJURP), as well as several accessory proteins and regulators (i.e. RSF1, MgcRacGAP, Condensin II, and KAT7) (Hu et al., 2011; Pan et al., 2017, 2019; Spiller et al., 2017). Although the deposition machinery is present throughout the cell cycle, new CENP-A deposition requires integrated signals from polo like kinase (PLK1) and cyclin-dependent kinases (CDK 1&2). PLK1 and CDK1/2 are differentially enriched across the cell cycle (Barr et al., 2004; Morgan, 1997). At the end of mitosis and early G1 phase, PLK1 phosphorylation of MIS18BP1 and HJURP complexes promotes their localization to centromeres, while high CDK1&2 kinase activity at all other timepoints phosphorylates and sequesters MIS18BP1 and HJURP away from centromeres (McKinley and Cheeseman, 2014; Müller et al., 2014; Pan et al., 2019; Silva et al., 2012; Spiller et al., 2017; Stankovic et al., 2017). Consistently, MIS18 complex formation, HJURP recruitment to centromeres, and CENP-A incorporation are all reduced in the presence of a small molecule PLK1 kinase inhibitor (McKinley and Cheeseman, 2014), whereas inhibition of CDK1 and CDK2 leads to ectopic deposition of CENP-A in G2 and S-phase (Silva et al., 2012; Stankovic et al., 2017). Importantly, dysregulation of any of these pathways or small changes in centromere composition renders the genome vulnerable to aneuploidy and genome instability, a hallmark of aging and many diseases including cancer (Barra and Fachinetti, 2018).

Although mechanisms governing CENP-A dynamics and inheritance in mitotic cells have been defined *in vitro*, our understanding of centromere regulation and maintenance *in vivo* remains limited (Prosée et al., 2020). Studies in *Drosophila* or plant species have suggested that CENP-A levels at centromeres are dynamically modulated during gametogenesis (Dattoli et al., 2020; Dunleavy et al., 2012; Raychaudhuri et al., 2012; Schubert et al., 2014). CENP-A levels in both the male and female germlines of these model organisms decrease transiently during meiosis but are subsequently restored in post-meiotic cells (Dattoli et al., 2020; Dunleavy et al., 2012; Raychaudhuri et al., 2012; Schubert et al., 2014). In contrast, CENP-A levels are not actively maintained, but are instead diluted in male and female meiotic and post-meiotic germ cells in mice (Das et al., 2022; Smoak et al., 2016). Indeed, a conditional knockout of the *Cenpa* gene in early prophase I-arrested mouse oocytes has no effect on centromere maintenance, oocyte maturation, or fertility (Smoak et al., 2016). These findings indicate that new CENP-A in mouse oocytes is not required for the extended period of oocyte quiescence, nor is additional CENP-A needed to resume meiotic divisions or support embryonic development (Smoak et al., 2016). Subsequent studies showed that heterozygous *Cenpa*^*+/-*^ in mothers decreases their live birth rate and reduces the absolute abundance of CENP-A protein at centromeres in the F1 offspring’s male germline and somatic cells (Das et al., 2022). These weakened centromeres are propagated trans-generationally, but interestingly, the epigenetic memory of reduced CENP-A in F1 mice is specifically erased in the female germline, providing F1 mothers with the CENP-A needed to support embryonic development of their offspring. Importantly, the epigenetic memory in the male germline and soma can be reset in 1-4 cell F1 embryos if the F0 *Cenpa*^+/-^ fathers are mated to wild type F0 *Cenpa*^+/+^ females (Das et al., 2022). Therefore, this correction process relies specifically on the maternal *Cenpa* genotype. Together, mammalian centromeres utilize genetic and epigenetic mechanisms to maintain and restore aberrations to centromeric chromatin from one generation to the next.

However, the physical inheritance and requirement of CENP-A at centromeres in the germline is variable across species (Mellone and Fachinetti, 2021). In the *Caenorhabditis elegans* germline, CENP-A is neither inherited nor required for re-establishment of the holocentric chromosomes in the early embryo (Gassmann et al., 2012). A similar transient absence of centromeric CenH3 has also been described in egg cells of *Arabidopsis thaliana*, which have regional CENP-A domains similar to mammalian centromeres (Ingouff et al., 2010). In *Drosophila melanogaster*, centromeric proteins such as CENP-C and Cal1, the ortholog of mammalian HJURP, are not retained in sperm (Dunleavy et al., 2012; Raychaudhuri et al., 2012). However, the CENP-A ortholog CID is essential for propagating the paternal genome to the next generation and generating a functional kinetochore during zygotic mitosis (Raychaudhuri et al., 2012). Specifically, embryonic inheritance of genetically tagged CENP-A nucleosomes from sperm is necessary for embryonic development and differing levels of paternally inherited CID are propagated into corresponding CID levels in the soma of offspring (Raychaudhuri et al., 2012). Case studies in humans have reported that the location and strength of neocentromeres, including those on the Y chromosome, can be passed down the paternal lineage (Amor et al., 2004; Bukvic et al., 1996; Tyler-Smith et al., 1999). In mice, somatic CENP-A levels are propagated faithfully from one generation to the next, but the extent and mechanism by which parental *Cenpa* transcripts and centromeric nucleosomes are required to re-establish functional centromeres in the totipotent embryo is unknown.

Here, by quantifying CENP-A dynamics throughout murine male and female gametogenesis, we find that CENP-A levels at centromeres are increased in germ cell precursors relative to somatic cells, but CENP-A loss is differentially regulated between the sexes, consequently leads to sperm retaining roughly a tenth of the CENP-A levels present in mature oocytes. Upon fertilization, chromatin-bound CENP-A levels begin to equalize between parental chromosomes. Unexpectedly, this equalization process includes physical redistribution of CENP-A from maternal pronucleus to the paternal genome, independent of zygotic transcription, translation, and DNA replication. Moreover, the mechanism that licenses re-localization of maternal CENP-A relies on mitotic centromere licensing factors: activation of PLK1 and inhibition of CDK1/2. Taken together, our data indicate that centromeres are reset from one generation to the next through active cooperation between the maternal and paternal pronuclei in the early embryo, despite their physical separation and differing epigenetic states. Furthermore, since this equalization process requires physical sharing of chromatin bound factors, our finding extends the window of opportunity for parental genomes to communicate, coordinate, and possibly compete.

## Results

### Generation and validation of *Cenpa*^*mScarlet/+*^ mice

To monitor CENP-A dynamics during mouse development and gametogenesis, we used CRISPR/Cas9 to tag the endogenous *Cenpa* gene locus with a flexible glycine-serine linker (GS), an enhanced red fluorescent protein mScarlet-I and a V5 epitope tag (Bindels et al., 2016; Hanke et al., 1992) (**Figure 1A**). We then confirmed the presence of the knock-in in two independent founder mice with Sanger sequencing (F0) and backcrossed these to C57Bl/6J mice for four generations. Overall, the *Cenpa*^*mScarlet/+*^ mice have no overt phenotypic differences or significant difference in testes/body weight ratio or mature sperm count relative to wild type litter mates (**Figures S1A&B**). Importantly, in cells of the seminiferous epithelium, the tagged CENP-A-mScarlet nucleosomes colocalized with CENP-C, a constitutive CCAN protein (**Figure 1B**), and defined anticentromere antibody (ACA) foci (**Figure S1C**). Furthermore, CENP-A-mScarlet puncta are exclusively localized to one end of the chromosomes in meiotic spreads of spermatocytes from *Cenpa*^*mScarlet/+*^ males, consistent with murine chromosomes being telocentric (Kalitsis et al., 2006) (**Figure S1D**). Taken together, the C-terminus tag on CENP-A does not appear to perturb CENP-A nucleosome function or localization.

**Figure 1:**
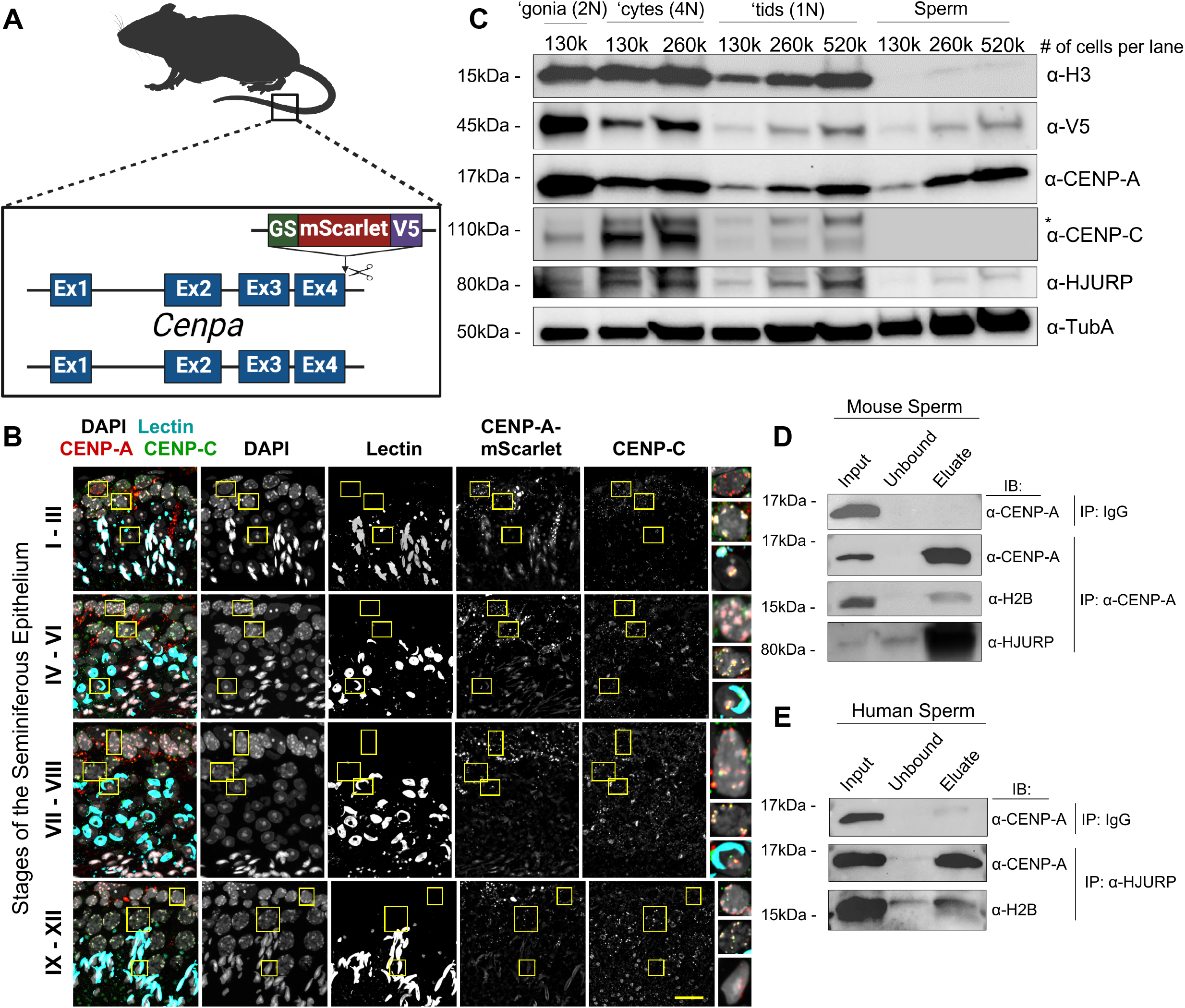
Reduction of CENP-A precedes the histone to protamine exchange in testes. A) Schematic of the Cenpa-GS-mScarlet-i-V5 transgene. B) Immunostaining of CENP-C in the *Cenpa*^*mScarlet/+*^ testis confirms CENP-C and tagged CENP-A colocalization across all stages and cell types within the seminiferous epithelium. Representative images from n = 3 staining experiments and n = 2 mice. Scale bar is 20um. C) Immunoblot of H3, V5, CENP-A, CENP-C, and HJURP in flow sorted spermatogonia (‘gonia), pachytene/diplotene spermatocytes (‘cytes), round spermatids (‘tids), and mature sperm. Alpha tubulin (TubA) is shown as a loading control. Shown is a representative immunoblot from n = 2 mice. Membrane was stripped and re-blotted for the proteins indicated. D) Immunoblot from CENP-A immunoprecipitation in mouse sperm, blotted for H2B and HJURP. Shown is a representative blot from n = 2 IPs on n = 2 mice. Membrane from the anti-CENP-A IP was stripped and re-blotted for the proteins indicated. E) Immunoblot from HJURP immunoprecipitation in human sperm, blotted for H2B and CENP-A. Representative blots from n = 3 HJURP immunoprecipitations on n = 3 deidentified human sperm samples. Membrane from anti-HJURP immunoprecipitation was stripped and re-blotted for H2B. IP = Immunoprecipitation, IB = Immunoblot.

### Reduction of CENP-A precedes the histone-to-protamine exchange in testes

During spermiogenesis, the male epigenome undergoes a near complete replacement of canonical histones with small basic proteins called protamines (Bao and Bedford, 2016), a process highly conserved across many vertebrates and invertebrates (Lewis et al., 2003). Although retention of CENP-A nucleosomes has been reported in mammals, invertebrates, and even plants (Dunleavy et al., 2012; Milks et al., 2009; Palmer et al., 1990; Raychaudhuri et al., 2012; Schubert et al., 2014), we do not know the extent to which CENP-A nucleosomes, like all canonical nucleosomes, are also replaced during spermiogenesis. To answer this question, we first enriched for germ cell populations isolated from adult *Cenpa*^*mScarlet/+*^ males using flow cytometry (Gaysinskaya et al., 2014). As expected, histone H3 levels are fairly constant between spermatogonia and meiotic spermatocytes, but decreases in post-meiotic haploid spermatids and is largely depleted in mature spermatozoa (**Figures 1C and S1E**). In contrast, the majority of tagged CENP-A-mScarlet is lost at the transition from spermatogonia to spermatocytes, and a small but notable loss occurs again during the transition from spermatocytes to round spermatids, with no significant change between spermatids and mature sperm (**Figures 1C and S1E**). We detected the same changes when assessing total CENP-A levels in germ cells at different stages (**Figures 1C and S1E**). Altogether, these data indicate that CENP-A levels are reduced during sperm development and that loss of CENP-A precedes the histone-to-protamine exchange.

### Centromere constituents change throughout spermatogenesis

Next, we examined the dynamics of centromeric proteins which directly interact with CENP-A. We found that CENP-C levels increase transiently in spermatocytes (with normalization for ploidy), despite significant loss of CENP-A, but decreases in haploid spermatids (**Figures 1C and S1E**). For CENP-B, which binds to B-box DNA sequences in minor satellites and interacts with CENP-A (Fachinetti et al., 2015; Fujita et al., 2015), we found that its levels were significantly reduced from spermatogonia to spermatocytes, but were maintained in spermatids (**Figures S1E&F**). For HJURP, the dedicated chaperone for CENP-A assembly, its protein levels were high in spermatogonia and spermatocytes but decreased in round spermatids. Finally, in mature sperm, both CENP-B and CENP-C were completely lost, consistent with earlier observations in Bull and *Drosophila* sperm (Palmer et al., 1990; Raychaudhuri et al., 2012), but to our surprise, low levels of CENP-A and HJURP persisted (**Figures 1C and S1E**). Furthermore, HJURP directly interacts or resides in the proximity of nucleosomal CENP-A and H2B, as evidenced by co-immunoprecipitation (co-IPs) of CENP-A and H2B from mouse sperm (**Figure 1D**) and reciprocal co-IPs from human sperm (**Figure 1E**). Taken together, these data indicate that centromeres are remodeled during germ cell differentiation, and that sperm-retained HJURP either associates with or localizes near to chromatin-bound CENP-A-containing nucleosomes, suggesting paternal HJURP may play a role in zygotic development.

### CENP-A differentially distributes between germ line stem cells and differentiating progenitor cells

In the gonad, the A_single_ (A_s_) spermatogonial cells undergo a series of mitotic divisions to give rise to undifferentiated spermatogonia (**Figure 2A**), consisting of syncytial clones of 2, 4, 8, and 16 cells interconnected by cytoplasmic bridges, referred to as A_paired_ (A_pr_), A_aligned-4_ (A_al-4_), A_al-8_, and A_al-16_. The A_s_, A_pr,_ and A_al_ cells are collectively referred to as A_undifferentiated_ (A_undiff_) spermatogonia, a heterogenous pool marked by two proteins: GFRα (A_s_, A_pr_, A_al-4_, and A_al-8_) and ID4 (A_s_ and A_pr_). The A_al_ differentiate into A1 spermatogonia, which undergo additional rounds of transit amplifying divisions to form A2, A3, A4, Intermediate, and Type-B spermatogonia (**Figure 2A**). Differentiating spermatogonia represent a transit amplifying progenitor pool marked by SOHLH1 and STRA8 expression that will undergo two meiotic divisions to generate haploid spermatids. To resolve CENP-A dynamics across the early stages of germ cell differentiation and compare relative CENP-A levels between germline and soma, we quantified fluorescence intensity in *Cenpa*^*mScarlet/+*^ intestinal villi and testis somatic cells (Sertoli & myoid) and compared these data to *Cenpa*^*mScarlet/+*^ levels in undifferentiated spermatogonia (GFRα1 and/or ID4) (Buageaw et al., 2005; Helsel et al., 2017), differentiating spermatogonia (SOHLH1 and STRA8) (Ballow et al., 2006; Endo et al., 2015; Zhou et al., 2008), and mid-meiotic prophase spermatocytes (γH2A.X) (Mahadevaiah et al., 2001) (**Figure 2A**). We found that the CENP-A-mScarlet fluorescence intensity was significantly higher in undifferentiated spermatogonia than any other gem cell subtype as well as intestinal villi epithelial cells and myoid cells in the testis (**Figures 2B&C**). Across germ cells, CENP-A-mScarlet intensity decreased in SOHLH1+ differentiating spermatogonia, even before the onset of meiosis. The reduction of CENP-A-mScarlet levels in differentiating spermatogonia cells (SOHLH1+ cells retain roughly 30% of somatic CENP-A-mScarlet levels), and the low levels maintained in Type-B spermatogonia and pachytene spermatocytes, consistent with the immunoblots in Figure 1C, indicate that the CENP-A reduction precedes early events of chromosome remodeling known to occur during meiotic prophase and that this reduction possibly links to germ cell differentiation. Unlike in flies and plants, where CENP-A levels decrease transiently during meiosis but are restored afterwards (Mellone et al., 2011; Schubert et al., 2014), mammalian spermatogenesis does not include a post-meiotic CENP-A recovery state, hence the mature mammalian spermatozoa are left with low levels of CENP-A (**Figure 1C**).

**Figure 2:**
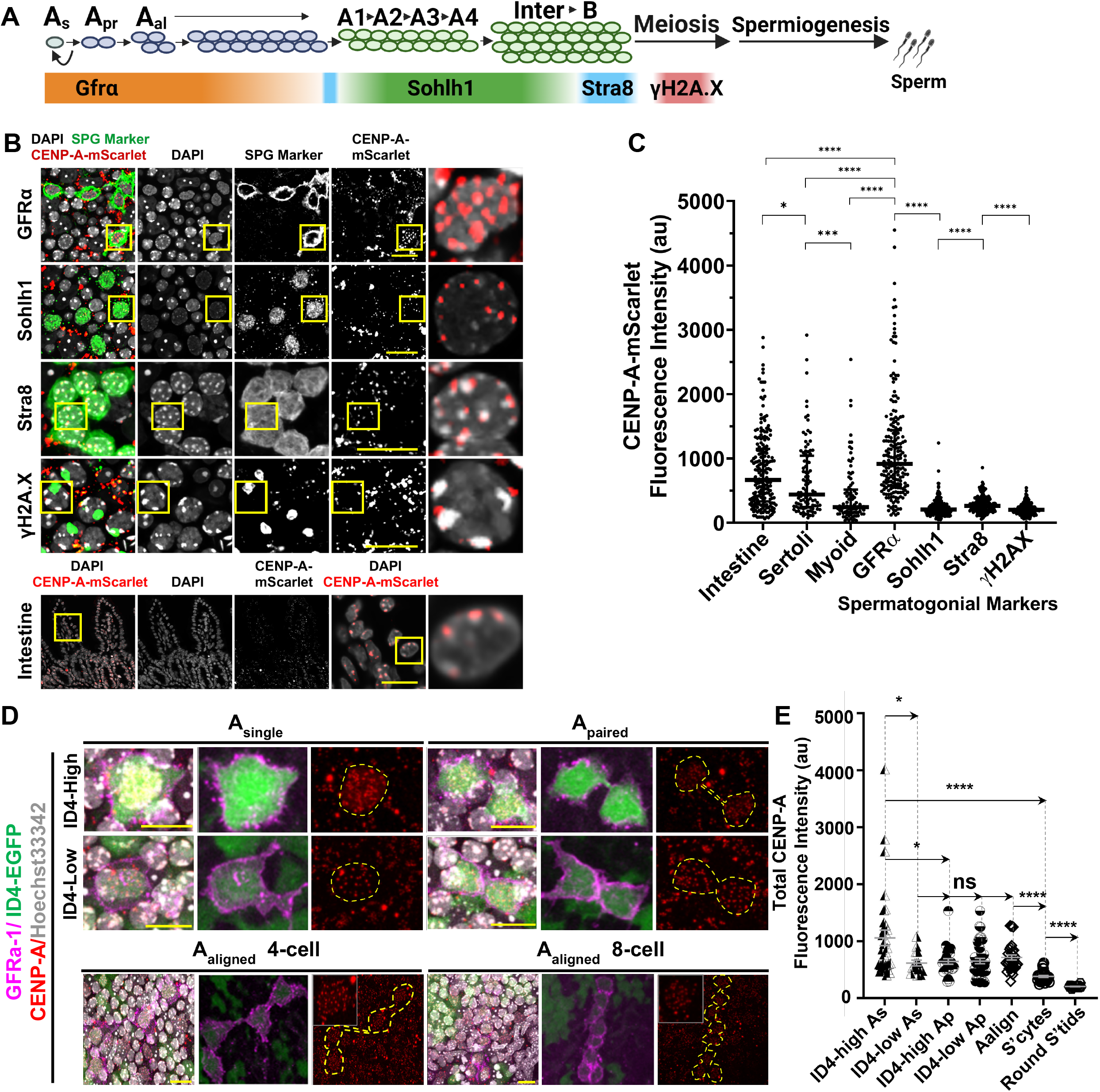
CENP-A asymmetrically segregates between germ line stem cells and differentiating progenitor cells. A) Schematic overview of the expression of indicated germ cell markers across spermatogenesis. Broadly, GFRα1 marks undifferentiated spermatogonia, SOHLH1 marks differentiating Type A and Intermediate spermatogonia, STRA8 marks differentiating Type B spermatogonia and early preleptotene spermatocytes, and condensed γH2A.X marks mid to late pachytene spermatocytes. B) Immunostaining of *Cenpa*^*mScarlet/+*^ whole mount tubules with the specified germ cell type markers. Representative images from n = 3 staining experiments on n = 3 mice. Scale bars are 20um. C) Quantification of CENP-A-mScarlet direct fluorescence across germ cell stages, somatic cells of the testes, and intestinal somatic cells. n = 200 germ and intestinal cells were quantified for each cell type, split evenly between n = 2 staining experiments and n = 2 mice. n = 100 Sertoli and Myoid cells were quantified for each cell type. Each dot is the sum of all CENP-A-mScarlet puncta in one cell. Mean fluorescence of CENP-A-mScarlet across the cell types are as follows: intestine = 668.5au, Sertoli = 441au, Myoid = 245au, GFRα = 918.5au, Sohlh1 = 206au, Stra8 = 264au, γH2A.X = 202.5au. D) Immunostaining of ID4-EGFP whole mount tubules with anti-GFRα1 showing different stages of undifferentiated spermatogonia. Representative images from n = 3 staining experiments on n = 4 mice. Scale bar is 10 µm. E) Quantification of total CENP-A immunofluorescence across all stages of ID4-High or ID4-Low undifferentiated spermatogonia, with spermatocytes (SPT) and round spermatids (RS) as controls. Mean fluorescence of CENP-A across the cell types is as follows: CENP-A_Id4 high As_ = 1059.53, CENP-A_Id4 low As_ = 617.45, CENP-A_Id4 high Ap_ = 645.07, CENP-A_Id4 high Ap_ = 653.15, CENP-A_Aalign_ = 645.42, CENP-A_SPT_ = 381.84, CENP-A_RS_ = 206.79. *: p < 0.05, **: p < 0.01, ***: p < 0.001, ****: p < 0.0001. ns indicates not significant.

To investigate potential biological underpinnings of the association between CENP-A reduction and spermatogonia differentiation, we leveraged the *Id4*^*eGFP/+*^ transgenic mouse line, in which functionally distinct A_s_ and A_pr_ spermatogonia cell populations can be distinguished based on their level of ID4 expression (Chan et al., 2014; Helsel et al., 2017; Sun et al., 2015). Among the hierarchical spermatogonial populations, the ID4-high A_s_ pool has the highest self-renewal capacity, as it generates the highest number of colonies after transplantation and forms the longest patches in lineage-tracing experiments (Helsel et al., 2017; Sun et al., 2015). In contrast, low ID4-expressing A_s_, A_pr_, and A_al_ cells have lower self-renewal capacity and transplantation efficiency, and are therefore believed to be transiently amplifying progenitors or differentiating spermatogonia (De Rooij, 2017). When we separated A_s_ and A_pr_ spermatogonia into ID4-High or ID4-Low fluorescence populations (**Figures 2D and S2A**) we found that the A_s_ population has higher ID4-eGFP fluorescence intensity than the A_pr_ spermatogonia, consistent with previous studies (**Figure S2B**) (Chan et al., 2014; Helsel et al., 2017). Quantification of total CENP-A immunofluorescence in spermatogonia revealed bulk differences in CENP-A levels: the ID4-High population has significantly higher levels of CENP-A than ID4-Low cells (**Figure S2C**). Interestingly, these population differences can be resolved further to reveal that the ID4-High A_S_ population has the highest levels of CENP-A among all staged germ cells, while ID4-Low A_s_, A_pr_, and A_al_ spermatogonia have 30% lower levels of CENP-A (**Figure 2E**). Thus, CENP-A loss occurs progressively throughout the differentiation trajectory of the male germline. The first reduction is observed at the stem cell to progenitor cell transition (A_s_ ID4-High to A_s_ ID4-Low), followed by subsequent rounds of reduction during transit-amplification and throughout meiosis.

Next, we asked whether the CENP-A distribution pattern in the mouse germline is conserved in *Drosophila melanogaster*. To this end, we collected adult fly testes and measured the total fluorescence intensity of the CENP-A ortholog CID within the somatic hub cells and in germ cell populations with increasing distance from the hub, representing various differentiation states: germline stem cells (GSCs), gonialblasts (GBs), spermatogonial cells (SGs, 4-16 cell stage) and primary spermatocytes (SCs, **Figure S2D**). Similar to our findings in the mouse germline, CID levels are increased in germline stem cells relative to the hub cells (**Figure S2F**), and gradually decrease during spermatogonial transit-amplification (**Figures S2E&F**); suggesting that CENP-A expansion and differentiation-coupled reduction is a conserved feature of male germ cell differentiation in both mice and flies.

### CENP-A density at centromeres increases from PGCs to GV oocytes

We then turned to investigate whether the CENP-A dynamics we observed in the male germline also occurs in the female germline during oogenesis. Unlike the continuous gametogenesis process in males, oogenesis in mice begins in the embryonic gonad (oogonia and preliminary stages of meiotic prophase I), followed by oocyte arrest at the dictyate stage of meiotic prophase I, a prolonged arrested phase which can last for the entirety of a female’s reproductive lifespan (Larose et al., 2019). To examine CENP-A and CENP-C dynamics in this protracted developmental process, we co-stained ovarian tissue cross-sections from Oct4^eGFP^ mice to quantify total CENP-A or CENP-C levels in primordial germ cells (PGCs) (E11.5) and in differentiating oogonia (E13.5, E18.5, P2, and adult) (**Figure S3A**) (Jones, 2008; Larose et al., 2019). As in the male germline, the early female germline progenitors, such as the PGCs and oogonia, have a higher level of CENP-A density at centromeres relative to the neighboring somatic cells, and the CENP-A levels at centromeres continue to increase between birth and adult germinal vesicle oocytes (GVs) (with normalization for ploidy) (**Figure S3B**). At the same developmental stages, CENP-C levels followed similar patterns to those seen for CENP-A: all assayed female germ cells contained more CENP-C than neighboring somatic cells, and the major increase in CENP-C occurred between birth and adult GV (**Figure S3C**). To monitor changes after meiotic resumption, we isolated GV-stage oocytes from *Cenpa*^*mScarlet/+*^, and wild type C57Bl/6J mice and *in vitro*-matured GV oocytes (4n) to generate metaphase I (MI) (4n) and metaphase II (MII) (2n) oocytes, as previously described (Stein and Schindler, 2011). We found that the elevated CENP-A density on oocyte centromeres are maintained in GV and MI, but decreases slightly between MI oocytes and MII eggs (after normalization for DNA ploidy) (**Figures S3D&E**). In contrast, CENP-C levels at centromeres steadily increased during female meiosis (normalized to DNA ploidy) (**Figure S3F**). Therefore, like males, CENP-A levels in early female germ cell precursors are significantly higher than what is reported in ovarian stromal cells, but unlike males CENP-A levels only slightly decrease between the MI to MII transition. Interestingly, in both male and female germlines, the decrease in CENP-A is concomitant with an increase in CENP-C, likely due to an active compensation mechanism for stabilization.

### Centromere epigenetic memory is cell-type and sex specific

Next, we examined how CENP-A levels in sperm compare to H3 levels in sperm or CENP-A levels in embryonic stem cells (ESCs) or oocytes. Consistent with earlier findings (Brinkley et al., 1986; Palmer et al., 1990; Sumner, 1987; Zalensky et al., 1993), we found a high concentration of CENP-A nucleosomes in the mature sperm of both mice and humans, relative to Histone H3 (**Figures S4A&B**). Despite the relatively higher retention, CENP-A levels in human and mouse sperm are roughly 12.5% or 17.5% of that in human and mouse ESCs respectively (**Figure S4C**). Curiously, although human sperm retains a higher overall percentage of Histone H3 nucleosomes than mouse sperm, mouse sperm retains more CENP-A nucleosomes than human sperm (**Figure S4C**). We speculate this difference in CENP-A retention may be due to differences in the amount of CENP-A needed to define centromere identity across species.

The striking difference in CENP-A levels between sperm and ESCs prompted us to ask how the levels in sperm compare to oocytes before uniting to generate a totipotent zygote, since unequal amounts of CENP-A levels in the gametes may pose a challenge for faithful chromosome segregation during zygotic divisions. In meiosis, chromosomes containing centromeres with unequal strengths are known to have biased segregation patterns between the oocyte vs. polar body (Chmátal et al., 2014; Iwata-Otsubo et al., 2017). Similarly in mitotic cells, a reduction in CENP-A levels results in chromosome segregation defects (Giunta and Funabiki, 2017; Li et al., 2011; Maehara et al., 2010). Therefore, functionally equivalent centromeres between homologous chromosomes must be critical for proper chromosome segregation. To compare the extent to which CENP-A levels differ between the two gametes, we collected and compared 450 GV and 450 MII eggs to a titration of sperm concentrations. Interestingly, after correcting for ploidy, CENP-A levels in sperm are significantly lower than in GV oocytes (∼17.5%) and MII eggs (∼10.6%) (**Figures S4D&E**). We confirmed that sperm do not retain CENP-C and that relative to GV oocytes, MII eggs carry close to twice the amount of CENP-C (183.7%) (**Figures S4D&E**). Interestingly, although CENP-A density on centromeres decreases, the total CENP-A pool, determined by immunoblots, increases from GV oocytes to MII eggs (170%), suggesting new CENP-A is translated during the final stages of oocyte maturation (**Figures S4D&E**) and could be essential for embryonic development (Smoak et al., 2016). Taken together, our results demonstrate that centromere remodeling/composition during gametogenesis is sex specific, leading to the generation of female and male gametes with epigenetically distinct centromeres in mouse.

### Paternal CENP-A-containing nucleosomes are inherited intergenerationally in mouse embryos

The requirement for intergenerational inheritance of centromeric nucleosomes varies across species (Das et al., 2017). To examine the fate of CENP-A nucleosomes in mouse sperm, we performed in-vitro fertilization (IVF) using MII eggs from superovulated C57Bl/6J females and sperm derived from *Cenpa*^*mScarlet/+*^ transgenic males. We collected embryos at various time-points in embryonic development (**Figure 3A**). Progression through the first embryonic cell cycle can be grouped into five pronuclear stages (PN1 to PN5) based on the size and relative location of the maternal and paternal pronuclei (Adenot et al., 1997; Wiekowski et al., 1997). Following fertilization, we visualized paternal CENP-A-mScarlet nucleosomes in the decondensing sperm head of PN0 stage embryos, throughout all pronuclear stages, and in the blastomeres of the 2-cell embryo after the first mitotic division (**Figure 3B**). Based on the presence of the tagged protein, we conclude that the paternal CENP-A nucleosomes in the embryo’s first cell cycle survive the protamine-to-histone exchange, DNA replication, and global genome reorganization. We only detected the CENP-A-mScarlet fluorescence signal on paternal chromatin, although it is possible that paternal CENP-A are removed and redistributed to the maternal pronucleus at a level undetectable by microscopy. To our knowledge, this is the first direct evidence of paternal nucleosome inheritance in the early mouse embryos.

**Figure 3:**
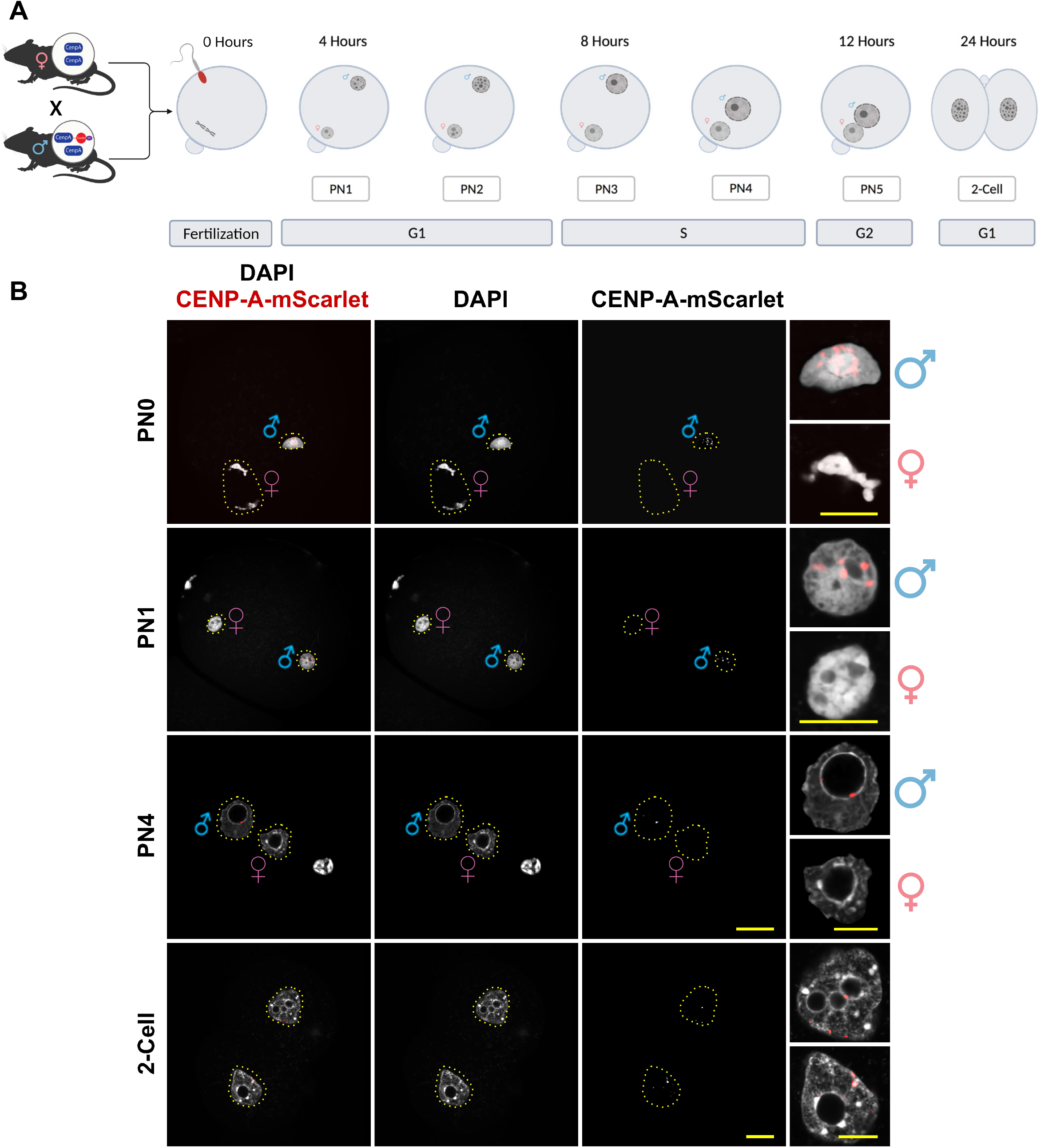
Paternal CENP-A-containing nucleosomes are inherited intergenerationally in mouse embryos. A) Overview of pronuclear staging in the first cell cycle after fertilization. Sperm is from *Cenpa*^*mScarlet/+*^ males and oocyte donors are C57Bl/6J. PN = pronuclear stage. B) Visualization of CENP-A-mScarlet fluorescence in *in vitro* fertilized embryos collected at the indicated pronuclear stage. Representative images from n = 4 IVF and immunostaining experiments using n = 4 males. Maternal and paternal pronuclei were identified by their relative size and position in relation to the polar body. Scale bars are 20um in main figure and 10um at the insets.

### Equalization of parental centromeres prior to zygote genome replication relies on maternally derived CENP-A

Next, we followed the dynamics and fate of maternally derived CENP-A nucleosomes. MII eggs were collected from superovulated *Cenpa*^*mScarlet/+*^ females and were *in vitro* fertilized using mature sperm from (C57Bl/6J X DBA2) F1 male mice (**Figure 4A**). As expected, maternal CENP-A-mScarlet nucleosomes were visible on condensed MII chromosomes and on chromosomes completing meiosis II in PN0 embryos (**Figure 4B**). The tagged CENP-A-mScarlet histones were observed throughout all pronuclear stages as well as in the blastomeres of two-cell embryos (**Figure 4B**). Unexpectedly, we found that the maternally inherited CENP-A-mScarlet histones began to localize to the paternal pronucleus at the end of PN2 and was found in both pronuclei of later staged zygotes (**Figure 4C**). This re-distribution of maternal CENP-A begins prior to the onset of zygotic S-phase in PN2-3 stage zygotes (**Figures 4C&D**) (Adenot et al., 1997; Wiekowski et al., 1997). By eight hours after fertilization (when most zygotes are in PN3), the majority of zygotes (77%) have maternal CENP-A-mScarlet in both pronuclei, with fewer having CENP-A-mScarlet in the maternal pronucleus (23%) (**Figure 4E**). Consistent with our earlier results measuring CENP-A levels by immunoblots, the quantification of CENP-A-mScarlet fluorescence in both pronuclei at fertilization (PN0-2; **Figure 4F**) showed that the sperm genome joins with roughly ∼6% of the maternal genome-derived CENP-A and that 80% of the CENP-A in the paternal pronucleus is inherited from the egg. Interestingly, when we measured total CENP-A levels in later staged embryos (PN5 zygotes), we found that centromeric chromatin was close to equal between parental pronuclei (paternal/maternal ratio: 0.87) (**Figure 4G**). These experiments suggest that equalization of CENP-A on centromeres is mostly complete before the first cell division and that the majority of CENP-A deposited on paternal centromeres is contributed by the maternal CENP-A.

**Figure 4:**
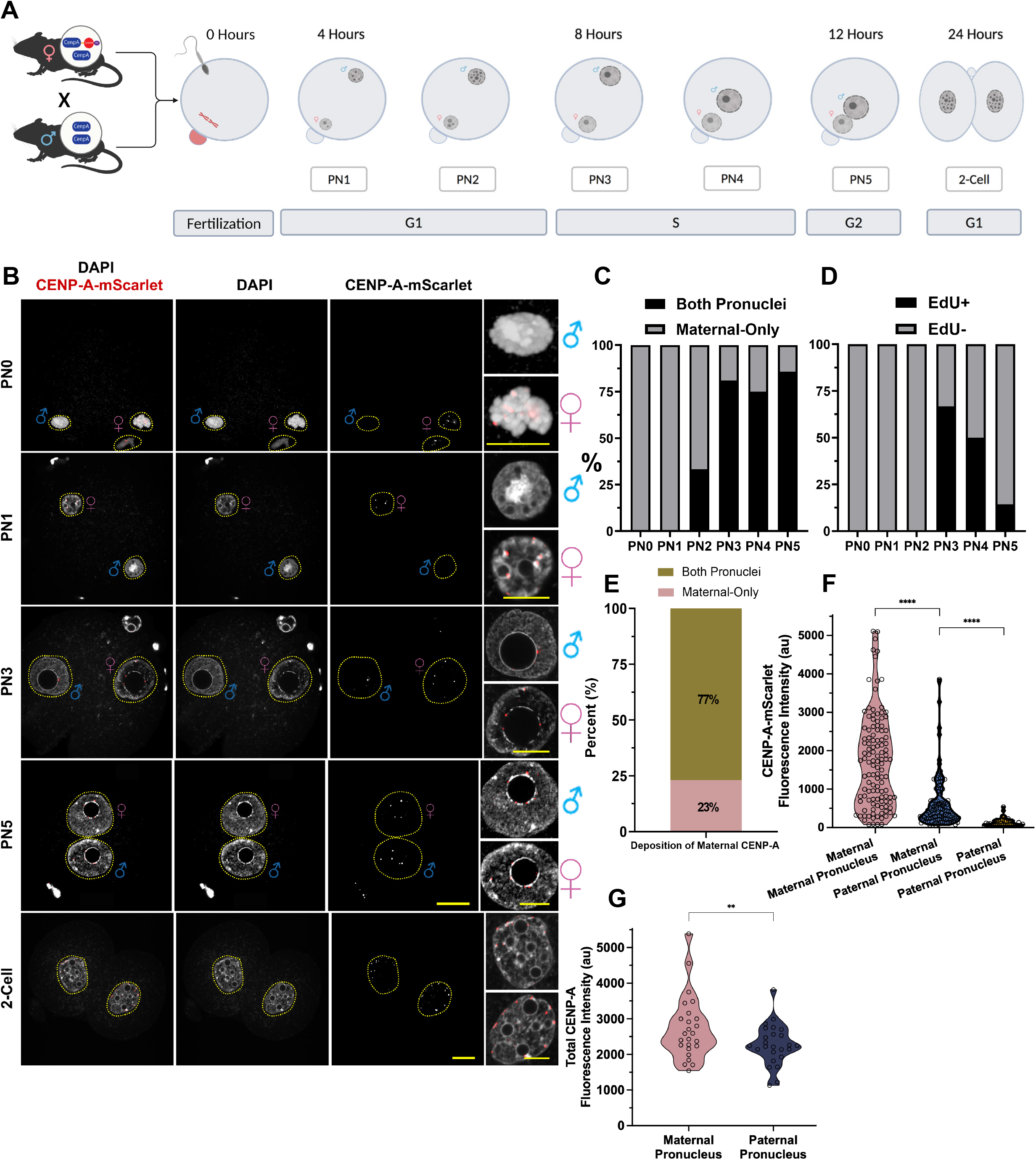
Equalization of parental centromeres prior to the first zygotic S-phase relies on maternally derived CENP-A histones. A) Overview of pronuclear staging in the first cell cycle after fertilization. Sperm is from C57Bl/6J males and oocyte donors are *Cenpa*^*mScarlet/+*^. B) Imaging endogenous maternal CENP-A-mScarlet fluorescence at the indicated pronuclear stages. Representative images from n = 4 IVF and immunostaining experiments using n = 12 females. Maternal and paternal pronuclei were identified by the relative pronuclear size and position in relation to the polar body. Scale bars are 20um in main figure and 10um at the insets. C) Quantification of the percentage of zygotes with maternal CENP-A-mScarlet in both pronuclei or only in the maternal pronuclei at each pronuclear stage. D) Quantification of the percentage of zygotes which stained positive or negative for EdU incorporation (indicating DNA replication). E) Quantification of the percentage of zygotes with maternal CENP-A-mScarlet found in both pronuclei or only the maternal pronuclei at 8 hours post fertilization. F) Comparing the direct fluorescence intensity of maternally vs. paternally derived CENP-A-mScarlet in either the male or female pronucleus at 8 hours post fertilization. Only zygotes with maternal CENP-A-mScarlet in both pronuclei were included, each dot is the sum of all CENP-A-mScarlet puncta in a single pronucleus. Mean fluorescence of maternal CENP-A-mScarlet in the maternal pronucleus = 1701au from n = 119 embryos. Mean fluorescence of maternal CENP-A-mScarlet in the paternal pronucleus = 416au from n = 103 embryos. Mean fluorescence of paternal CENP-A-mScarlet in the paternal pronucleus = 103au from n = 35 embryos. ****: p < 0.0001. G) Quantification of total CENP-A immunofluorescence in the maternal and paternal pronuclei at stage PN5, each dot is the sum of all total CENP-A puncta in a single pronucleus. Mean fluorescence of total CENP-A in the maternal pronucleus = 2583au from n = 25 embryos. Mean fluorescence of total CENP-A in the paternal pronucleus = 2245au from n=25 embryos. **: p < 0.01. PN = pronuclear stage

### Equalization of parental centromere density is independent of zygotic transcription, translation, and DNA replication

Oocytes contain a significant reservoir of mRNA and protein that supports embryonic development prior to ZGA (Gao et al., 2017; Tadros and Lipshitz, 2009; Yartseva and Giraldez, 2015) but supposedly do not carry a large pool of free cytosolic CENP-A proteins (Smoak et al., 2016). Next, we examined whether transcription of either *Cenpa-mScarlet* mRNA or satellite repeats during the minor ZGA wave is necessary for deposition of maternal CENP-A-mScarlet onto paternal centromeres. To this end, we collected MII eggs from super ovulated *Cenpa*^*mScarlet/+*^ females and performed IVF using mature sperm from (C57Bl/6J X DBA2) F1 male mice. At 1.5 hours after fertilization, we treated zygotes with 200μM 5,6-dichloro-1-β-D-ribofuranosyl-benzimidazole (DRB), a reversible RNA polymerase II inhibitor known to inhibit the minor ZGA wave in mice (Abe et al., 2018), or vehicle control (DMSO). To monitor transcription in the presence or absence of the inhibitor, we incubated embryos with 500nM 5-ethynyl-uridine (EU), a uridine analog that can be detected using click chemistry reactions (Jao and Salic, 2008), and collected embryos either 10 hours after fertilization or at the 2-cell stage (24-30hrs) (**Figures 5A and S5G**). As expected, the DRB treatment inhibited transcription in 1- and 2-cell embryos, as determined by total EU signals in vehicle and DRB-treated embryos (**Figures 5A and S5G**, note that EU staining has cytoplasmic background in all assayed cell types and treatments). Interestingly, inhibition of the minor ZGA wave did not prevent maternal CENP-A-mScarlet from being redistributed to the paternal pronucleus (**Figures 5A&D**). Quantification of maternal CENP-A-mScarlet revealed no significant differences in CENP-A-mScarlet intensity between the DRB-treated maternal and paternal pronuclei relative to the control (**Figures 5A&D**). The drug treatment also did not significantly alter cell cycle progression (**Figure S5D**). Altogether, these results indicate that neither transcription of *Cenpa* or transcription-mediated nucleosome turnover are necessary for maternal CENP-A-mScarlet to be localized to paternal genome.

**Figure 5:**
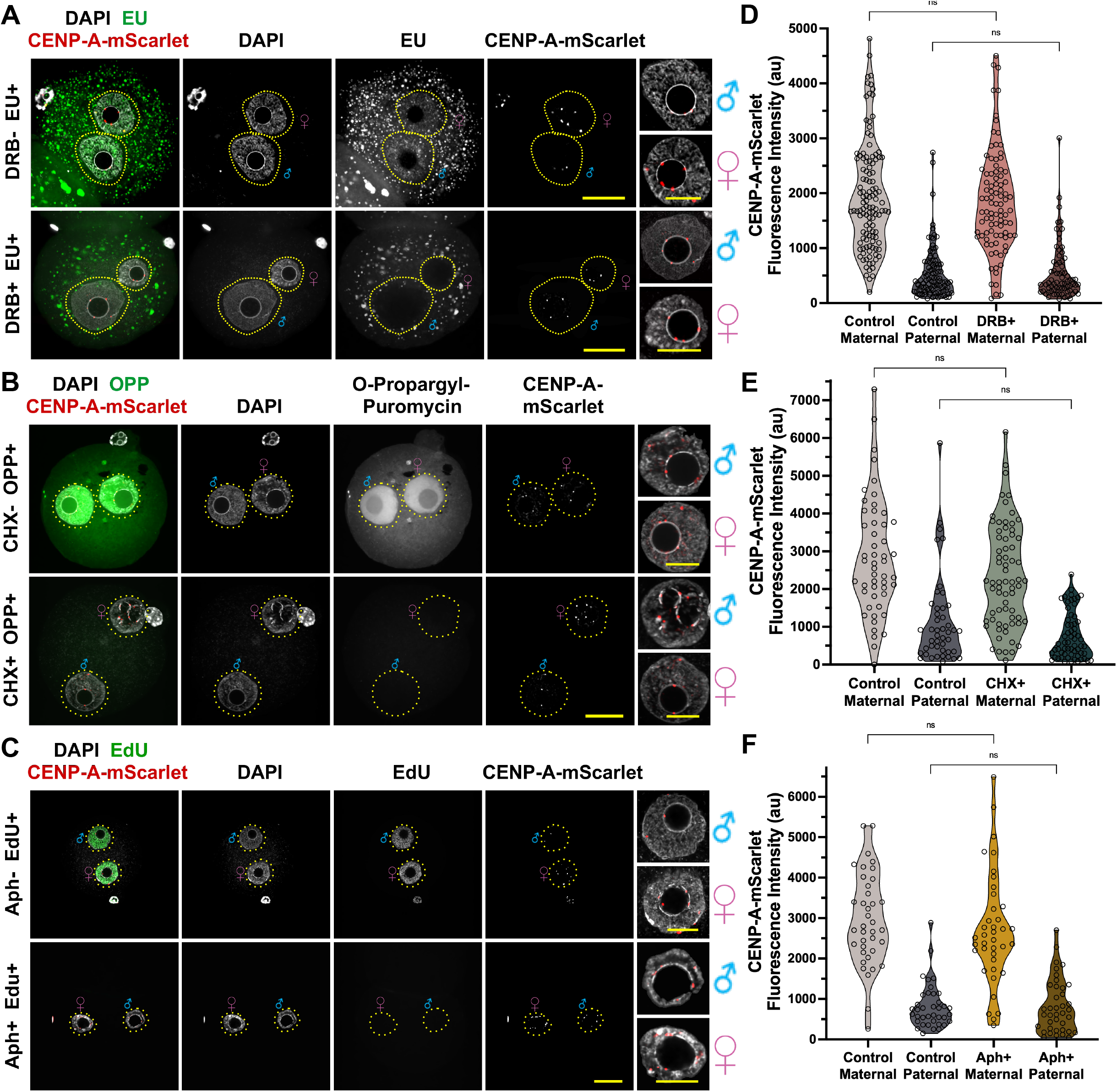
Equalization of parental centromere density is independent of zygotic transcription, translation, and DNA replication. A-C) Representative images of zygotes collected from: A) Embryos treated with 200uM 5,6-dichloro-1-β-D-ribofuranosylbenzimidazole (DRB). Note 5-ethynyl uridine (EU) is used to monitor RNA synthesis and drug treatment efficiency. Embryos were collected at 10hpf from n = 5 IVFs with n = 13 *Cenpa*^*mScarlet/+*^ females. B) Embryos treated with 100ug/mL cycloheximide (CHX). Note: O-propargyl puromycin (OPP) is added to the culture media to monitor nascent peptide synthesis. Embryos were collected at 8.5hpf from n = 3 IVF experiments with n = 9 *Cenpa*^*mScarlet/+*^ females. C) Embryos treated with 10ug/mL Aphidicoline (Aph). Note 5-ethynyl deoxyuridine (EdU) is used to monitor DNA synthesis. Embryos were collected at 8.5hpf from n = 3 IVFs with n = 9 *Cenpa*^*mScarlet/+*^ females. Scale bars are 20um in main figure and 10um at the insets. D-F) Quantification of CENP-A-mScarlet direct fluorescence from embryos treated with DRB (D), CHX (E), Aph (F). Each dot is the sum of CENP-A-mScarlet puncta in each pronucleus. Mean fluorescence intensities shown in D) are as follows: Control maternal = 1741au from n = 119 embryos, Control paternal = 390.5au from n = 98 embryos, DRB+ maternal = 1688au from n = 91 embryos, DRB+ paternal = 359au from n = 71 embryos. Mean fluorescence intensities shown in E) are as follows: Control maternal = 2849au from n = 51 embryos, Control paternal = 856au from n = 45 embryos, CHX+ maternal = 2348au from n = 69 embryos, CHX+ paternal = 551au from n = 61 embryos. Mean fluorescence intensities shown in F) are as follows: Control maternal = 2952au from n = 38 embryos, Control paternal = 681.5au from n = 38 embryos, Aph+ maternal = 2708au from n = 39 embryos, Aph+ paternal = 650au from n = 39 embryos. ns indicates not significant.

Next, we investigated whether translation of oocyte-derived *Cenpa* mRNA was responsible for the increase in CENP-A observed in the paternal pronucleus. When we used RT-qPCR with Taqman probes for *Cenpa* across different stages, we found that GV oocytes carry higher levels of *Cenpa* mRNA relative to mESCs. However, *Cenpa* mRNA decreases during meiotic maturation and fertilization, and then steadily increases in late-zygotes, 2-cell, and 4-cell embryos, before finally dropping sharply in 8-cell embryos (**Figure S4F**). These measurements indicate that oocytes do provide *Cenpa* mRNA which can be translated and used as a potential source of CENP-A protein in the paternal pronucleus. To test this possibility, we treated IVF-derived embryos with either 100ug/mL cycloheximide (CHX) or an equal volume of vehicle (DMSO) after 1 hour of fertilization and cultured the zygotes for approximately another seven hours. To verify that the CHX treatment was effective, we added O-propargyl-puromycin (OPP), which incorporates into the nascently synthesized polypeptide chains and can be stained by click chemistry as a read out for translational activity (Liu et al., 2012). We confirmed that the OPP signal was absent in CHX treated embryos (**Figure 5B**). Interestingly, inhibition of protein synthesis did not significantly change the amount of maternal CENP-A-mScarlet levels being deposited into the paternal pronucleus (**Figures 5E and S5B**), beyond a statistically insignificant decrease between control and CHX-treated pronuclei (**Figure S5E**). To eliminate potential bias from cell cycle differences, we re-analyzed only PN3 embryos. We still did not detect any significant reduction of CENP-A-mScarlet in the male pronuclei. However, we found a slight, but statistically significant decrease in the maternal pronuclei of CHX-treated embryos (**Figure S5H**). These results suggest that post-transcriptional control of *Cenpa* mRNA in the zygote helps establish steady state CENP-A levels at least in the maternal pronucleus. Curiously, blocking translation does not appear to significantly affect CENP-A deposition into the paternal pronucleus. Although translation of *Cenpa* mRNA contributes to maternal CENP-A levels in the early embryo, its impact is small (a 29% decrease in maternal pronuclei CENP-A due to CHX-treatment). Clearly, further study is needed to understand the origin of maternal CENP-A deposited in the paternal pronucleus.

Finally, since the majority of embryos have maternal CENP-A-mScarlet nucleosomes in both pronuclei near the time of zygotic S-phase, we assessed whether DNA replication is necessary for the re-localization of maternal CENP-A-mScarlet into the paternal genome. To this end, we followed a similar IVF schema, but incubated zygotes with 10ug/mL aphidicolin (Aph), a reversible DNA polymerase inhibitor that blocks DNA synthesis (Ikegami et al., 1978) or vehicle (DMSO), starting 1.5 hours after fertilization, and assessed incorporation of 5-Ethynyl-2′-deoxyuridine (EdU), a thymidine analog, to confirm the inhibition of DNA synthesis (**Figure 5C**). Interestingly, inhibition of zygotic DNA replication did not prevent maternal CENP-A-mScarlet from being redistributed to the paternal pronucleus (**Figures 5C&F**), and the drug treatment did not drastically alter the pronuclear staging of Aph-treated embryos (**Figures S5F**). Taken together, these experiments suggest that paternal incorporation of maternal CENP-A-mScarlet does not require embryonic transcription, translation, or DNA replication, although some contribution from *de novo* translation is possible.

### Equalization of parental centromere densities relies on the redistribution of maternally derived nuclear CENP-A

Since the levels of newly translated and deposited CENP-A-mScarlet represent a minority of total centromeric CENP-A, we asked whether pre-existing, chromatin-bound maternal CENP-A nucleosomes could be a source of the maternal CENP-A re-distribution to the paternal genome. To test this possibility, we collected *Cenpa*^*mScarlet/+*^ GV oocytes, photobleached mScarlet fluorescence only in the nuclei, and performed oocyte *in vitro* maturation, followed by *in vitro* fertilization to monitor the levels and dynamics of CENP-A-mScarlet and total CENP-A (**Figure 6A**). Our photobleaching protocol depleted ∼80% of the CENP-A-mScarlet signal in the nuclei of GV oocytes while maintaining oocyte integrity and competency (**Figures 6B&C; see methods**). In live cell cultures, photobleaching of *Cenpa*^*mScarlet/+*^ GV oocytes caused a fluorescence signal intensity drop from an average of 3115au to an average of 252au (**Figures 6B&C**) that recovered to 665au an hour after bleaching but did not significantly increase after another 24 hours of recovery (**Figures 6B&C**). Given these robust responses, we subsequently performed IVF on *in vitro*-matured oocytes that were or were not photobleached, cultured the embryos in the presence of 100ug/mL cycloheximide to eliminate any contribution from new translation, and collected embryos eight hours after fertilization (note: equalization is complete, see **Figure 6A**). We found that CENP-A-mScarlet signal intensity was significantly decreased in both the maternal and paternal pronuclei, resulting in a 68.5% and 37% decrease in fluorescence, respectively (**Figures 6D&E**); suggesting that photobleached CENP-A-mScarlet in the nuclei of GV oocytes is re-localized to the paternal chromosomes of early embryos. Importantly, the total CENP-A levels measured by immunofluorescence did not significantly change between photobleached and control embryos (**Figure 6F**), indicating that equalization is occurring despite the decrease in CENP-A-mScarlet fluorescence. After accounting for incomplete photobleaching, we estimate that roughly 50% of CENP-A-mScarlet in the paternal pronucleus and ∼85% in the maternal pronucleus originates from the nuclei of GV oocytes. In these photobleaching experiments we find a measurable level of CENP-A-mScarlet in both pronuclei and the amount of CENP-A in the female pronucleus appears higher than the levels retained after photobleaching or extended culture. This may be due to more photorecovery occurring after fertilization than during GV oocyte culture, and we are thus monitoring the dynamics of photo recovered CENP-A. More likely, both the maternal and paternal centromeres increase their CENP-A density using a limited pool of pre-existing cytoplasmic and/or the newly translated oocyte CENP-A observed in Figure S4D&E. Taken together, we conclude that equalization and expansion of both parental centromeres relies on two sources: 1) CENP-A-mScarlet from the GV oocyte’s nucleus being re-localized to the paternal chromatin of early embryos, and 2) some deposition of a maternal pool of translated CENP-A in both parental pronuclei.

**Figure 6:**
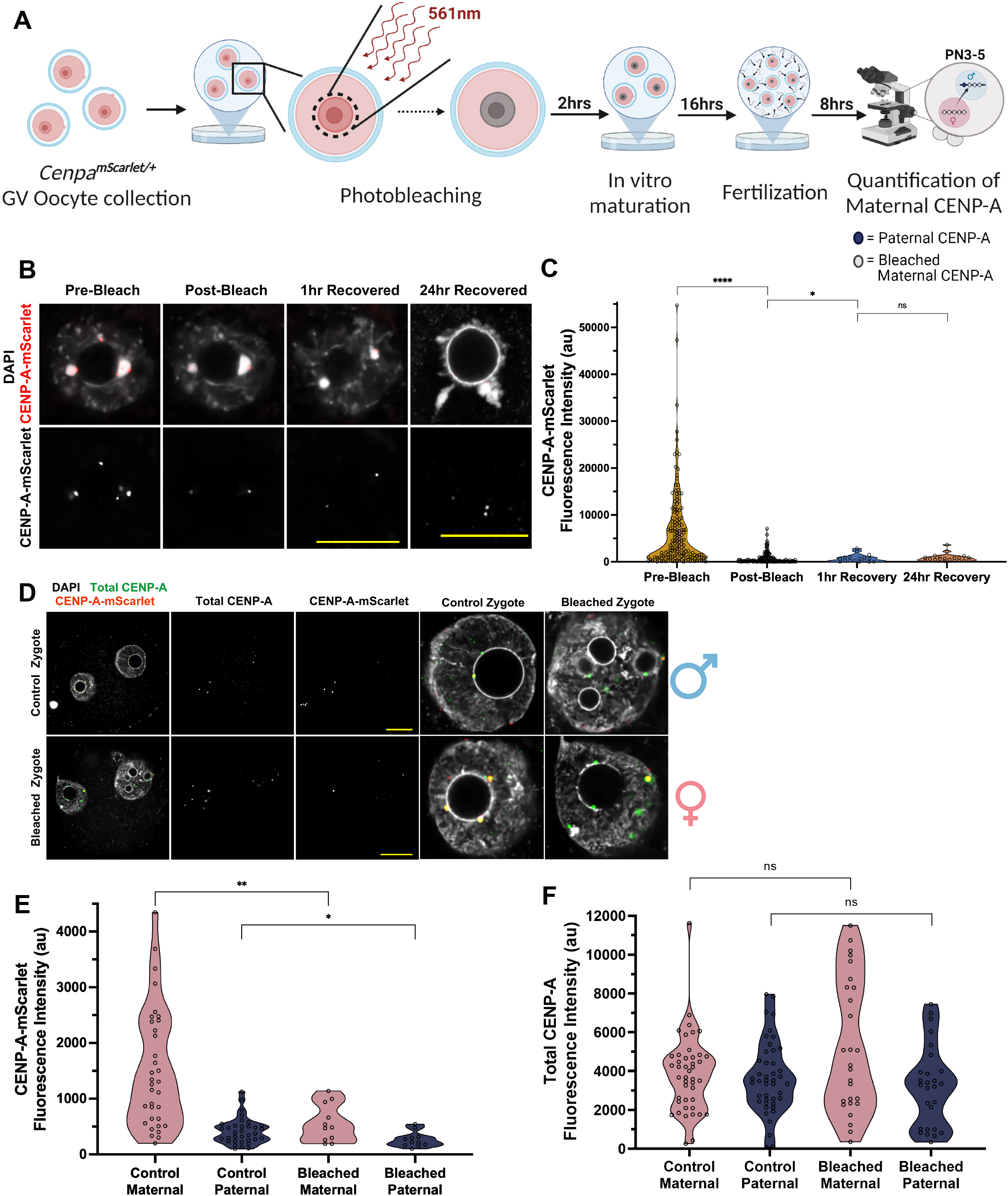
Equalization of parental centromere densities relies on the redistribution of maternally derived nuclear CENP-A. A) Overview of maternal CENP-A-mScarlet photobleaching experiments. Sperm is from (C57Bl/6J X DBA2)F1 males and oocyte donors are *Cenpa*^*mScarlet/+*^. B) Representative images of CENP-A-mScarlet endogenous fluorescence in oocytes prior to photobleaching, immediately post photobleaching, and after recovering for 1hr or 24hrs. C) Quantification of CENP-A-mScarlet direct fluorescence in oocytes pre-bleach, immediately post bleach, and 1 and 24 hours after recovery. Each dot is the sum of CENP-A-mScarlet puncta in one oocyte. Mean fluorescence intensities shown in C) are as follows: Pre-bleach = 3115au from n = 161 oocytes, post-bleach = 252au from n = 161 oocytes, 1hr recovery = 665au from n = 17 oocytes, 24hr recovery = 668.5au from n = 20 oocytes. D) Representative images of CENP-A-mScarlet endogenous and total CENP-A fluorescence in PN3 zygotes from either a control or photobleached experiment. All zygotes were incubated with 100ug/mL cycloheximide as in Fig. 5B. Images are representative of n = 5 independent photobleaching experiments using a total of 10 *Cenpa*^*mScarlet/+*^ females. All scale bars are 20um. E&F) Quantification of either direct CENP-A-mScarlet fluorescence (E) or total CENP-A immunofluorescence (F) from the PN3-PN5 embryos represented by panel D. Each dot is the sum of CENP-A-mScarlet or total CENP-A puncta in one pronucleus. Mean fluorescence intensities shown in E) are as follows: Control maternal = 1467au from n = 41 embryos, Control paternal = 401au from n = 41 embryos, bleached maternal = 476au from n = 16 embryos, bleached paternal = 253au from n = 16 embryos. Mean fluorescence intensities shown in F) are as follows: Control maternal = 3205au from n = 46 embryos, Control paternal = 2738au from n = 46 embryos, bleached maternal = 2276au from n = 30 embryos, bleached paternal = 1838au from n = 30 embryos. *: p < 0.05, **: p < 0.01, ****: p < 0.0001. ns indicates not significant.

### CENP-A levels in zygotes are predetermined by the available pool of CENP-A and deposition machinery

The female germline is a striking example of tightly controlled centromere maintenance. Oocytes arrested in prophase I of meiosis show little to no exchange of chromatin-bound CENP-A (see 24hr FRAP in **Figures 6B&C**). A previous study that injected *Cenpa* RNA into oocytes demonstrated that CENP-A deposited in the prenatal ovary is sufficient to support embryonic development until at least ZGA, and that CENP-A nucleosomes stably assembled on centromeres persisted for more than a year after genetic ablation (Smoak et al., 2016). More recently, heterozygous *Cenpa*^*+/-*^ mice were used to track the epigenetic memory of reduced CENP-A levels intergenerationally (Das et al., 2022). In this study, the maternal genotype of the *Cenpa* locus was found to be important for resetting centromeric CENP-A levels in zygotes and impacted reproductive fitness and litter size. For example, if oocytes derived from *Cenpa*^*+/-*^ heterozygous mice are fertilized by sperm with decreased CENP-A, then the low CENP-A levels will be epigenetically maintained on paternal chromosomes from the single cell embryo into the adult offspring. However, if the mother is *Cenpa*^*+/+*^, weak paternal centromeres will have their CENP-A levels equalized to wild-type maternal levels by the 4-cell embryonic stage (Das et al., 2022).

Here, to further explore the link between cytoplasmic CENP-A in oocytes and zygotic CENP-A deposition, we overexpressed tagged CENP-A in GV oocytes and tracked tagged CENP-A in the early embryo. To this end, we microinjected *Cenpa-Emerald* RNA into GV oocytes isolated from wild type CF1 female mice at 6-10 weeks of age. After 2-4 hours of recovery and translation of the CENP-A-Emerald protein, we *in vitro* matured the oocytes and either sampled stages of oocyte maturation (GV, MI, and MII stages) or fertilized injected eggs with (C57Bl/6J X DBA2) F1 sperm. We then collected the resulting embryos at 8.5 hours after fertilization. Although the injected *Cenpa-Emerald* RNA was efficiently translated in GV oocytes, and cytoplasmic protein was easily visualized in all subsequent stages of meiotic female germ cells, we did not observe any centromeric localization of the injected CENP-A-Emerald (**Figure S6A**), in agreement with previously published results (Smoak et al., 2016). Instead, CENP-A-Emerald was deposited at ectopic genomic loci in MI and MII eggs (**Figure S6A**), a euchromatic mis-localization that persisted after fertilization (**Figure S6A, PN0/3**), and is likely mediated by the promiscuous association between the Histone H3.3 chaperones DAXX or HIRA and free CENP-A/H4 dimers (Lacoste et al., 2014; Nye et al., 2018).

As we observed in our CENP-A-mScarlet transgenic lines, CENP-A-Emerald injected into oocytes consistently became deposited at the centromeres of PN3 and later stage zygotes (**Figure S6A**, compare PN0/PN3 insets). When we quantified the total CENP-A immunofluorescence at centromeres of zygotes 8.5 hours after fertilization, we found a significant increase of total CENP-A levels in both maternal and paternal pronuclei relative to non-injected controls (**Figure S6B**). We also found that cytoplasmic CENP-A-Emerald is preferentially loaded onto the centromeres of the paternal pronucleus (**Figures S6A&B**). These results confirm that CENP-A inherited from the cytoplasm of oocytes can function as a source for equalizing CENP-A levels across maternal and paternal centromeres and indicates that a sufficient excess of inherited RNA and protein can expand CENP-A levels beyond the original genetic background. Interestingly, the forced increase in CENP-A-Emerald deposition begins in PN3 or later stage embryos; indicating that oocytes and early zygotes utilize a centromere licensing mechanism to prevent maternal CENP-A deposition until the late G1 phase (PN2-3) in zygotes.

Next, we tested whether co-injecting CENP-A and the HJURP deposition machinery could further increase CENP-A deposition. Indeed, we found that the co-injection significantly increased total CENP-A levels in the female pronucleus relative to un-injected controls and *Cenpa-Emerald*-only injected embryos (**Figure S6B**). Surprisingly, we found no significant difference between paternal pronuclei (**Figure S6B**). These results suggest that maternally inherited HJURP is not a limiting factor for CENP-A deposition in the paternal pronucleus, and that HJURP retained in mature sperm may play a functional role in guiding paternal CENP-A deposition in the zygote (**Figures 1C-E**). Altogether, these experiments support a mechanism in which embryonic CENP-A levels are determined by the inherited pool of maternal CENP-A protein, and equalization of centromeric chromatin in early embryos occurs through an asymmetric deposition mechanism which favors maternal CENP-A to be deposited onto the paternal genome.

### Asymmetric deposition of CENP-A is correlated with asymmetric levels of CENP-C and MIS18BP1

Tight regulation of the localization and abundance of the CENP-A deposition machinery ensures that a single wave of centromeric nucleosomes are deposited during late telophase and early G1 phase of the cell cycle (Müller and Almouzni, 2017). In mammalian mitotic cells, CENP-C associates with CENP-A throughout the cell cycle and directs the MIS18 complex and HJURP to restore CENP-A levels in G1 (Falk et al., 2015; Guo et al., 2017; Mitra et al., 2020). To assess localization patterns in embryogenesis, we generated wild-type zygotes by *in vitro* fertilizing C57Bl/6J oocytes with (C57Bl/6J X DBA2) F1 sperm and collected embryos at staggered timepoints across the first cell cycle. Despite absence of CCAN proteins CENP-B or CENP-C in sperm (**Figures 1C and S1E&F**), we found that maternal CENP-C is deposited immediately into the decondensing sperm genome in PN0-stage embryos, preceding CENP-A, and localizes to centromeres at all stages of the first embryonic cell cycle (**Figure S7A**). Quantification of CENP-C revealed that paternal centromeres load more CENP-C than maternal centromeres, and that centromeric CENP-C levels are reduced during zygotic development (**Figures S7A&B**). In addition to the increased CENP-C enrichment in the paternal pronucleus, we found that MIS18BP1 localizes to pronuclei by PN2 and becomes enriched in the paternal pronucleus of PN2-3 zygotes before being gradually lost in later stage embryos (**Figure S7C**). MIS18BP1 is also colocalized with puncta of maternal CENP-A-mScarlet on the paternal genome (**Figure S7E, arrowheads**). Together, these results suggest that the accumulation of centromere deposition and maintenance factors prior to CENP-A deposition is a conserved feature between mitotic and zygotic centromeres and may contribute to the equalization of parental centromeres. The paternal enrichment of both proteins also provides a possible explanation for the asymmetric deposition of overexpressed CENP-A-Emerald into paternal centromeres: preferential loading of constitutive centromere associated proteins at sperm chromatin after fertilization could bias subsequent recruitment of the centromere deposition machinery in a self-propagating manner.

### CENP-A redistribution in the zygote is licensed by CDK and PLK1 kinase activities

In cultured human cells, small molecule inhibition of CDK1/2 is well known to induce ectopic CENP-A deposition during the S and G2 phases, while inhibition of PLK1 in mitotic cells prevents CENP-A deposition regardless of CDK activity (McKinley and Cheeseman, 2014; Müller et al., 2014; Pan et al., 2019; Silva et al., 2012; Spiller et al., 2017; Stankovic et al., 2017). In the mouse embryos, the presence of the pan-CDK inhibitor Flavopiridol modulates the cell cycle to cause only 80% of 1-cell embryos to progress into the 2-cell stage (Oqani et al., 2011), whereas the presence of the small molecule and selective PLK1 inhibitor BI2536 does not impact zygotic development until the metaphase-to-anaphase transition (Baran et al., 2013). Here, we set out to investigate the role of CDK1 and PLK1 kinases during the equalization of parental CENP-A in early embryos.

First, we titrated each drug to determine the effective inhibitor concentration in zygotes and found that 5uM of Flavopiridol and 10nM BI2536 were sufficient to allow >80% of treated zygotes to develop to the 2-cell stage (data not shown). Then, we collected MII eggs from super-ovulated *Cenpa*^*mScarlet/+*^ females, performed IVF using mature sperm from (C57Bl/6J X DBA2) F1 male mice, treated zygotes 1.5 hours after fertilization with either 5uM Flavopiridol or vehicle control (DMSO) and collected samples 8.5 hours after fertilization. Although levels of CENP-A-mScarlet in the maternal pronucleus were not impacted by CDK inhibition, the amount of CENP-A-mScarlet deposited in the paternal pronucleus increased (**Figures 7A&C**). The drug treatment did slow the cell cycle progression in Flavo-treated embryos (**Figure S7E**), and also increased the fraction of PN2 embryos with CENP-A-mScarlet deposited in both pronuclei (compare **Figure S7F and Figure 4C, Figure S7H**). These data suggest that CDK1/2 activity contributes to the timing and amount of CENP-A deposition in the paternal pronucleus of the zygote. Next, we treated zygotes with 10nM BI2536 and found that inhibition of PLK1 kinase activity significantly decreased CENP-A-mScarlet levels in both the maternal and paternal pronucleus (**Figures 7B&D**). BI2536 treatment slowed cell cycle progression (**Figure S7G**), and reduced the percent of zygotes with maternal CENP-A in both pronuclei (**Figure S7H**). Since inhibition of PLK1 reduces maternal CENP-A-mScarlet levels to that of MII eggs and paternal CENP-A to levels slightly higher than that of sperm (**Figure 7D**), we conclude that PLK1 activity is required for CENP-A deposition in the maternal pronucleus and contributes to the expansion of CENP-A in the male pronucleus. Taken together, these experiments suggest that zygotes and mitotic somatic cells utilize similar mechanisms to license centromeric deposition. Furthermore, the differential contribution of PLK1 and CDK1/2 activity to CENP-A expansion in the maternal vs paternal pronuclei suggest that our previous findings of asymmetric CENP-A deposition (**Figure S6B**) in zygotes may be due to differences in the mechanism of deposition or licensing in the two parental pronuclei.

**Figure 7:**
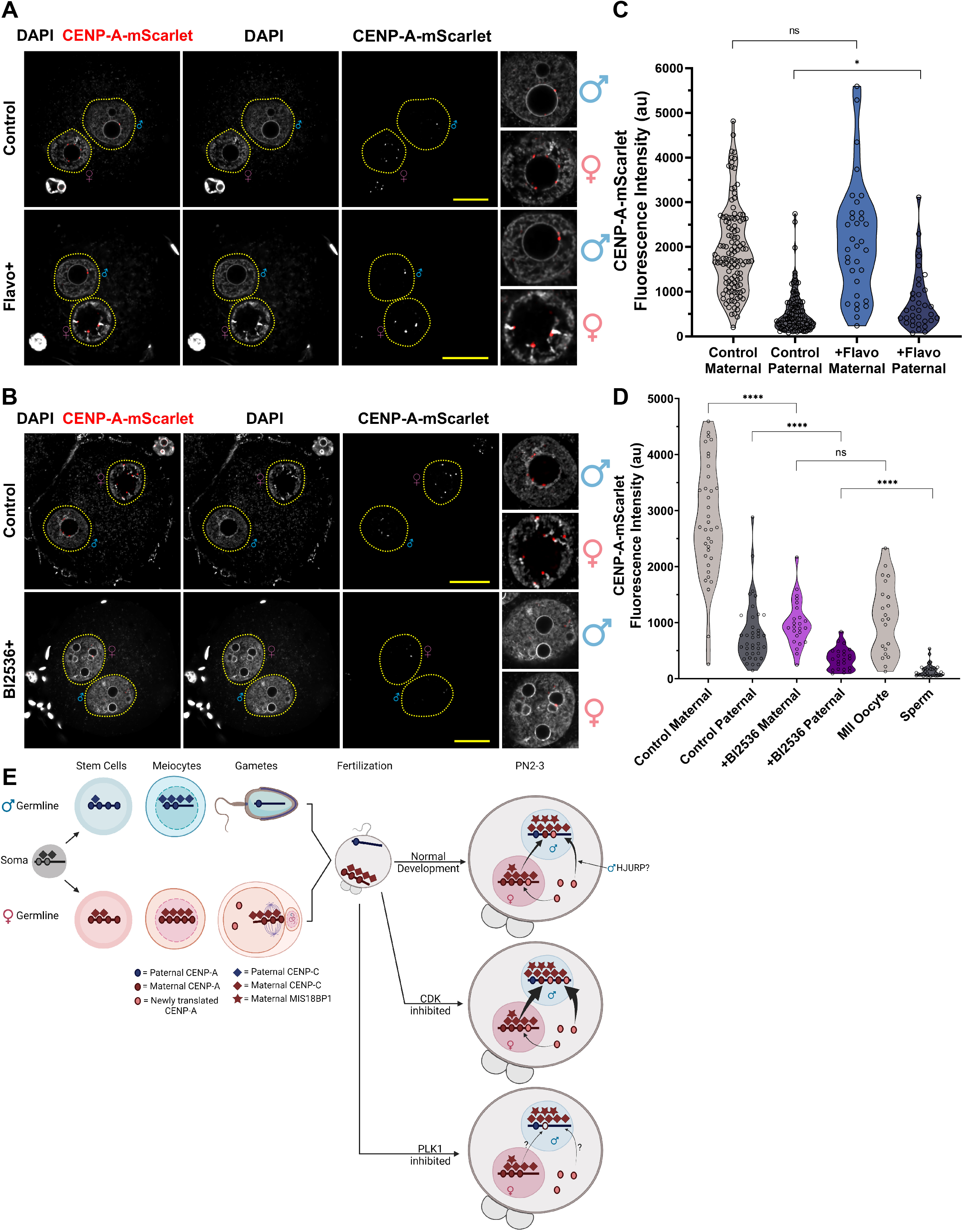
CENP-A redistribution in the zygote is licensed by CDK and PLK1 kinase activity. A&B) Representative images of zygotes collected from IVF experiments as in Fig 5. A) 5uM Flavopiridol (Flavo) treatment. Embryos were collected at 8.5hpf from n = 2 IVF experiments with n = 4 *Cenpa*^*mScarlet/+*^ females. B) Embryos treated with 10nM Bl2536. Zygotes were collected at 8.5hpf from n = 2 IVFs with n = 4 *Cenpa*^*mScarlet/+*^ females. Scale bars are 20um or 10um in insets. C&D) Quantification of CENP-A-mScarlet direct fluorescence from embryos treated with Flavo (C) or Bl2536 (D). Each dot is the sum of CENP-A-mScarlet puncta in each pronucleus. Mean fluorescence intensities shown in C) are as follows: Control maternal = 2348au from n = 69 embryos, Control paternal = 551au from n = 61 embryos, Flavo+ maternal = 2849au from n = 51 embryos, Flavo+ paternal = 856au from n = 45 embryos. *: p < 0.05, ns indicates not significant. Mean fluorescence intensities shown in D) are as follows: Control maternal = 2703au from n = 36 embryos, Control paternal = 681.5au from n = 36 embryos, BI2536+ maternal = 917.5au from n = 24 embryos, BI2536+ paternal = 351au from n = 24 embryos, MII eggs = 1086au from n = 21 oocytes, and sperm (paternally inherited CENP-A-mScarlet, see **Figure 4F**) = 103au from n = 35 embryos. ****: p < 0.0001, ns indicates not significant. E) Model of CENP-A maintenance in germ cells and early embryos.

## Discussion

Among all chromosomal features, the centromere is perhaps most unique because it is solely defined epigenetically by the histone H3 variant CENP-A. The centromere is also pivotal for proper cell division due to its central organizing roles of recruiting kinetochores and anchoring spindle microtubules. However, our current knowledge about centromeres is limited since their underlying maintenance and regulation mechanisms are very context-dependent. Accumulating evidence suggests that different cell types and cells from different developmental stages often display varied characteristics and employ distinct mechanisms to propagate functional centromeres and epigenetic memory. By following CENP-A dynamics across the mouse germline and soma, we found that CENP-A levels are much higher in germline stem cells than in somatic cells. Although similar dynamics are conserved between mice and flies, the relative levels of CENP-A at centromeres in germline stem cells, differentiating cells, and somatic cells differs across species, reflecting cell-type and species-specific differences in CENP-A maintenance programs. The differential regulation of CENP-A density at centromeres may be one mechanism to establish tissue and sex specific differences in centromere identity, but in a subset of flies and mosquito species, differences in centromeric states in the germline and soma can be achieved by using distinct CENP-A paralogs, indicating that distinct mechanisms are employed evolutionarily to achieve tissue and cell-type specific composition and structure (Finseth et al., 2015; Kawabe et al., 2006; Kursel and Malik, 2017; Kursel et al., 2020; Maheshwari et al., 2015). Variation in centromere density and composition leads us to speculate how such differences in centromere packaging are achieved. We hypothesize that CENP-A post-translational modifications (PTMs) or cell type-specific centromere-associated factors can modify centromeric chromatin composition and structure. Indeed, many CENP-A PTMs have been identified, but their functions are either unexplored or currently under debate, and many of the PTMs identified in humans are not conserved even within the mammalian clade (Srivastava and Foltz, 2018). A comprehensive assessment of CENP-A PTMS and interactors across cell type may be needed to understand cell type-specific regulation and centromere homeostasis.

Although centromere expansion is a unique property of germline stem cells, differences in CENP-A and centromere maintenance mechanisms have been observed in quiescent progenitors and terminally differentiated and nondividing cells (Milagre et al., 2020; Swartz et al., 2019). Our data in the female germline suggest that CENP-A levels in oocytes increase during early stages of folliculogenesis: from primordial oocytes in P2 to adult GV oocytes (**Figure S3B**), but CENP-A levels at centromeres do not seem to increase from the GV stage and onwards, even when ectopic *Cenpa* RNA or *Cenpa* + *Hjurp* is injected into GV oocytes (**Figures S6A&B**). Rather, our data show that CENP-A levels at centromeres are diluted across MI-MII divisions. The increase in CENP-A density at centromeres from primordial to GV oocytes appears to be inconsistent with earlier studies (Smoak et al., 2016) which suggested that CENP-A levels at centromeres do not change in GV oocytes when comparing wild-type and *Gdf9-Cre;Cenpa*^*fl/fl*^ mouse lines, despite losing *Cenpa* mRNA transcripts from P2 onwards in the *Gdf9-Cre;Cenpa*^*fl/fl*^ line. We think these differences may be due to the timing of actual *Gdf9-Cre* mediated excision of the *Cenpa* gene as well as translation of a trace pool of *Cenpa* mRNA. However, the increase or resetting of CENP-A density we observe in the maternal germline between primordial to GV oocytes explains the mechanism and timing of the erasure of centromere memory observed in *Cenpa*^+/-^ female germline (Das et al., 2022).

In males, CENP-A proteins levels are differentially regulated. The CENP-A levels are tightly maintained across self-renewing progenitors, but are significantly decreased upon the commitment to differentiation (**Figure 2E**). The decreased CENP-A in differentiating progenitors suggests that the asymmetric CENP-A distribution may result from an asymmetric division of A_single_ cells. Whether the CENP-A asymmetry dictates spermatogonial cell fate or if CENP-A levels serve as biomarker for the fate of the spermatogonial cells are intriguing questions and will be an active area of further investigation. Asymmetric segregation of H3 and H4 (old vs. new) as well as and CENP-A is a hallmark of asymmetric cell division identified in *Drosophila* male germline stem cells (Ranjan et al., 2019; Tran et al., 2012) and in *Drosophila* intestinal stem cells (Chen and Zion, 2020; García del Arco et al., 2018). CENP-A was also found to be inherited asymmetrically in female germline stem cells (Dattoli et al., 2020). Moreover, *in vitro* CENP-A levels can modulate cell fate, as CENP-A overexpression can promote quiescence vs. epithelial mesenchymal transition (EMT) in cells depending on P53 status (Jeffery et al., 2021), underscoring a potential link between centromeric protein levels, genome reprogramming, and cell fate. Taken together, the expansion of mammalian germline centromeres may provide a mechanism to maintain functional centromeres throughout differentiation despite the lack of any significant CENP-A deposition events between the two meiotic divisions.

Curiously, the sex-specific differences in CENP-A regulation at centromeres leads to the generation of sperm and egg with varied centromeric strengths, signifying a new epigenetically imprinted embryonic asymmetry in a parent-of-origin-specific manner. In addition to CENP-A, maternal and paternal gametes bring or acquire asymmetric amounts of CENP-B, MIS18BP1, CENP-C, and HJURP in early embryos, which may underscore the asymmetric deposition of CENP-A between the paternal and the maternal pronucleus. In male and female germline stem cells in *Drosophila* the CENP-A chaperone chromosome alignment defect 1 (CAL1 in male) (Ranjan et al., 2019) or the inner kinetochore component CENP-C (in female) (Carty et al., 2021) have been found to be required for the deposition of CENP-A and maintenance of the CENP-A asymmetry between the two daughter cells derived from stem cell asymmetric divisions. Notably, it has been reported that dual spindles form in early mammalian embryos to separate parental genomes, and they have been proposed to underlie the error-prone cleavages in early mammalian embryos (Reichmann et al., 2017). Whether this dual spindle has evolved to adapt to the centromeric differences of two gametes is unclear. Nevertheless, given the central roles of centromeres in mitosis, this novel equalizing process identified in mammalian zygotes could serve as an elegant mechanism to prevent detrimental chromosomal segregation defects. Therefore, our work has broader implications for understanding centromere maintenance, inheritance, and segregation as biological processes that safeguards normal development and may help us better understand the causes of genomic instability observed in many cancers and early embryonic development.

Importantly, paternal CENP-A nucleosomes in sperm are inherited and maintained until the 2-cell stage and are sufficient to define and re-establish centromeres in paternal chromosomes in the zygote. Although many studies have utilized indirect methods to suggest that paternal nucleosomes are inherited and can affect early embryonic development (Amor et al., 2004; van der Heijden et al., 2008; Tyler-Smith et al., 1999; Van De Werken et al., 2014), to our knowledge, our study is the first physical demonstration and direct visualization of intergenerational epigenetic inheritance. Conceivably, our observations of CENP-A marked nucleosomes may extend to other types of histones retained in sperm, suggesting that paternal nucleosomes can serve as epigenetic memory carriers from father to offspring. It is also possible that CENP-A inheritance mechanisms are unique, as these nucleosomes are more stable across mitotic divisions than H3.1 or H3.3-marked nucleosomes (Bodor et al., 2013; Falk et al., 2015; Hemmerich et al., 2008). Additionally, their inheritance may also be contingent on continued association of CENP-A with its chaperone HJURP in sperm. Indeed, HJURP binding to CENP-A nucleosomes during passage of the replication fork maintains centromeric localization during mitotic divisions (Huang et al., 2015; Nechemia-Arbely et al., 2019; Zasadzińska et al., 2018). We hypothesize that HJURP mediates a similar stabilizing effect on CENP-A during the global chromatin remodeling process (protamine to histone exchange) that occurs in the zygotes after fertilization.

Recently, it has been shown that both the maternal *Cenpa* genotype and the centromeric DNA sequences contribute to zygotic centromere strength (Das et al., 2022). In this work, mouse models with compromised CENP-A levels (*Cenpa*^*+/-*^) or genetically different mouse strains with unequal centromeric satellite sequences were used to generate hybrid progeny to study both the maternal and genetic contribution to centromere propagation. In contrast, we used genetically identical parents in our studies. Therefore, the difference reported here is purely epigenetic and due to the dimorphic gametogenesis between sexes. Technically, in their studies CENP-A immunofluorescent intensities were measured and compared without distinguishing CENP-A in a parent-of-origin-specific manner. Contrastingly, here we used genome editing to endogenously tag the *Cenpa* gene and performed a series of *in vitro* fertilization approaches to directly visualize the transfer of maternally derived CENP-A from the maternal pronucleus to the paternal pronucleus. To our knowledge, this is the first time such an epigenetic equalization process has been visualized in mammalian embryos *in vivo*. Finally, we found that equalizing CENP-A between the maternal pronucleus and the paternal pronucleus requires PLK1 activity, while CDK1/2 only affects CENP-A expansion in the paternal pronucleus but has little effect on the maternal pronucleus. Therefore, our studies reveal a novel and asymmetric epigenetically imprinted phenomenon at centromeres which utilizes chaperones and mitotic licensing factors to equalize paternal differences in early mammalian embryos.

Altogether, our results uncover novel insights into the regulation of centromeric chromatin in the mammalian germline and early embryos. Expansion of CENP-A levels in spermatogonia and oogonia allows the stem cells to retain functional centromeres across multiple mitotic and meiotic divisions, which each result in a minimal loss of CENP-A. We propose a model of CENP-A maintenance in early embryos where CDK1/2 initially inhibits CENP-A deposition on paternal chromosomes in the early zygote. Centromere licensing factors such as CENP-C and MIS18BP1 can then enrich asymmetrically onto paternal chromatin and direct preferential CENP-A deposition into the paternal chromatin prior to the entry of the zygotic S-phase in a PLK1-dependent manner. While the precise mechanism which directs this asymmetric loading of the CENP-A deposition machinery is unknown, paternal HJURP inherited from sperm seems a parsimonious candidate. Therefore, wild-type oocytes can utilize both cytoplasmic and re-distributed nuclear maternal CENP-A to equalize parental centromeres and erase the epigenetic memory of paternal centromeres with low levels of CENP-A (**Figure 7E**).

## Acknowledgements

We thank Drs. Richard Schultz, Paula Stein, and Carmen Williams, for their invaluable scientific discussions on *in vitro* maturation of oocytes and embryos, and the Hammoud Laboratory members for discussions. We also thank Dr. Haiqing Zhao for his continuous support of the project and for co-mentoring B.M., and Jon Oatley for sharing the Id4-eGFP mouse line. This work wouldn’t be possible without the assistance of Dr. Thomas Saunders in designing targeting vectors and members of the University of Michigan Transgenic Animal Model Core in the generation of the transgenic mouse model. Portions of Figures 6 and 7 were created with BioRender.com. This research was supported by National Institute of Health (NIH) grants 1DP2HD091949-01 (S.S.H.), R01HD104680 01 (S.S.H), R01HD058730 (A.D., M.A.L., B.E.B.), F31HD100124 (G.M.), R35GM127075 (X.C), R35GM136340 (M. A., K.S.), training grants T32GM145470 (G.M.) and T32GM007544 (C.T.), Rackham Predoctoral Fellowship (G.M.), and Open Philanthropy Grant 2019-199327 (5384) (S.S.H.). X.C. is a Howard Hughes Medical Institute Investigator.

## Author contributions

G. M. and S.S.H. conceived and designed the experiments. G.M. carried out majority of experiments with some help from K.J., B.M., C.T., S.C. M.A., and R.R. G.M. and S.S.H. received experimental and model system expertise from K.S. and X.C. respectively. G. M., C.T., and S.S.H. wrote the manuscript and received input from all co-authors especially from K.S. and X.C. S.S.H supervised the project.

## METHODS

### EXPERIMENTAL MODEL AND SUBJECT DETAILS

#### Mice

All experiments utilizing animals in this study were approved by the Institutional Animal Care and Use Committees of the University of Michigan (Protocols: PRO00006047, PRO00008135, PRO00010000), John Hopkins University (Protocol: MO22A71), and Rutgers University (Protocol: 201702497) and was performed in accordance with the National Institutes of Health Guide for the care and use of laboratory animals. Briefly, mice were housed in an environment controlled for light (12 hours on/off) and temperature (21 to 23C) with *ad libitum* access to water and food (Lab Diet #5008 for breeding mice, #5LOD for non-breeding animals).

*Cenpa*^*mScarlet/+*^ knock-in mice were generated on the C57BL/6N background using CRISPR/Cas9-mediated genome editing by the University of Michigan’s Transgenic Animal Model Core. The sgRNA and donor oligo were designed as previously described (Haeussler et al., 2016). The guide RNA target sequence was selected according to the on- and off-target scores provided by the web tool CRISPOR (Haeussler et al., 2016) (http://crispor.tefor.net) and proximity to the target site. Ribonucleoprotein (RNP) complexes were formed by mixing the sgRNA (2.5 ng/uL) with Cas9 protein (IDT, 5 ng/uL) in Opti-MEM (ThermoFisher) and incubating at 37C for 10 minutes, at which time the donor oligo (IDT, 10 ng/uL) containing the intended transgene was added. Zygotes from super-ovulated C57BL/6N females were microinjected into the paternal pronucleus with the RNP/donor oligo mix using a Femtojet 4i microinjector (Eppendorf). After injection, zygotes were moved to KSOM AA medium (Sigma), matured to the 2-cell stage, and transferred to oviductal ampullas of pseudopregnant CD-1 females. All animal procedures were carried out in accordance with the Institutional Animal Care and Use Committee and approved protocol of the University of Michigan. Offspring were genotyped for the GS-mScarlet-I-V5 transgene insertion by extraction of genomic DNA from a small ear biopsy. Mouse lines from two founding CRISPR/Cas9 transgene insertions were maintained separately and utilized interchangeably in all experiments. Transgenic males and control mice were used for all experiments at 8-16 weeks of age.

#### Collection and preparation of human sperm

The University of Michigan Institutional Review Board determined this study did not fit the definition of research involving human subjects (U.S. Department of Health & Human Services regulations at 45CFR46.102) because the research was intended to contribute to generalizable knowledge, the researchers did not interact with human subjects, nor obtained identifiable private information or identifiable biospecimens.

Discarded and de-identified semen samples were obtained from the University of Michigan Center for Reproductive Medicine. Briefly, semen samples were collected via masturbation from men presenting for routine semen analysis. Samples were allowed to liquefy at room temperature in a sterile container for 30 minutes before being processed for storage. The total semen samples were washed three times in PBS with centrifugation at 200g to remove seminal fluid and somatic cells in the resulting pellet were lysed in PBS (Thermo Fisher) + 0.1% SDS (Sigma) + 0.5% Triton-X-100 (Sigma) for 30 minutes on ice. The samples were centrifuged at 200g for 10 minutes and the final sperm pellet was snap frozen in liquid nitrogen and stored at − 80C.

#### Culture of human and mouse embryonic stem cells

Human and mouse embryonic stem cell (ESC) lines were used in this study. All protocols for the use of the human ESC lines were approved by the Human Pluripotent Stem Cell Research Oversight Committee at the University of Michigan. The human H1 (WA01, P32) male ESC line was purchased from WiCell (NIHhESC-10-0043, Lot #RB18522) and cultured in Sue O’Shea’s lab at the University of Michigan Pluripotent Stem Cell Core. Briefly, the H1 ESCs were grown in feeder-free conditions in 60mm petri dishes (Thermo Fisher) coated with Matrigel (BD Biosciences) and maintained in mTESR1 media (Stem Cell Technologies) at 37C with 5% CO2. H1 cells were checked daily for differentiation and passaged every 4 days using Dispase solution (Thermo Fisher). The mouse E14TG2a male ESC line originated from the European Collection of Authenticated Cell Cultures (Sigma 08021401, Lot #15H010). The E14TG2a ESCs were cultured in gelatin-coated (Sigma) 6-well culture plates (Thermo Fisher) in Glasgow’s MEM media (Thermo Fisher) supplemented with 50 units/mL Penicillin-Streptomycin (Thermo Fisher), 0.1 mM Nonessential amino acid (Thermo Fisher), 1 mM sodium pyruvate (Sigma), 0.1 mM β-mercaptoethanol (Thermo Fisher), 1000 units/mL Leukemia Inhibitory Factor (Sigma), and 10% fetal bovine serum (Thermo Fisher). The mESCs were maintained at 37C with 5% CO2, checked daily for differentiation, and passaged at 80% confluency with TrypLE (Thermo Fisher). All cell lines were karyotypically normal and negative for mycoplasma at the indicated passage.

## METHOD DETAILS

### Phenotypic assessment of *Cenpa*^*mScarlet/+*^ males

All phenotyping was carried out in males between 8 and 10 weeks of age. All weight measurements were recorded less than 5 minutes after euthanasia. Sperm were counted using a Makler chamber (Cooper Surgical SEF-MAKL) and performed as n=3 independent technical replicates per mouse (n=3 mice per founding line).

### Microscopy

All samples were imaged on a Nikon A1R HD25 point scanning confocal microscope with a 60X 1.4NA oil objective (Nikon MRD71600) and motorized stage (Nikon TI2-S-SE-E) with 405nm, 488nm, 561nm, and 64nm solid state laser lines detected using a 1-channel GaAsP spectral detector (Nikon A1 DUV-B HD). The microscope platform was controlled in Nikon NIS Elements (Version 5.21.03). Images were acquired with 0.75um z-steps and ∼140nm xy pixel resolution.

### Immunofluorescence on cryosectioned mouse testes

Testes were collected from wild-type C57BL/6J or *Cenpa*^*mScarlet/+*^ males between 8-12 weeks of age. Mice were euthanized by cervical dislocation and their testes were immediately removed and snap frozen in liquid nitrogen. The frozen testes were then embedded in OCT medium (Leica 39475237) on dry ice and stored at − 80C in a sealed container. After embedding, the tissue was cryosectioned onto glass coverslips (Thermo Fisher) at a thickness of 10um and stored at −80C. To prepare for staining the coverslips were removed from storage at −80C and dried for 10-20 minutes at room temperature. The tissue sections were then fixed in 4% paraformaldehyde (Millipore Sigma) for 10 minutes, washed in PBS (Thermo Fisher) two times for 5 minutes each, and permeabilized in PBS + 0.5% TritonX-100 (Thermo Fisher) for 30 minutes at room temperature. Next, the tissue sections were washed twice in PBS, then blocked for 1 hour at room temperature in PBS + 3% BSA + 0.1% TritonX-100 filtered through a 0.2um filter (Millipore Sigma) before incubation with primary antibodies diluted in PBS + 3% BSA + 0.1% TritonX-100 filtered through a 0.2um filter at 4C overnight: rabbit α CENP-A at 1:250 dilution (Cell Signalling Technologies 2048), rabbit α CENP-C at 1:2000 dilution (gift from Ben E. Black). The next day, the tissue sections were washed four times for 15 minutes each at room temperature in PBS + 0.1% TritonX-100 then labelled with PNA Lectin-FITC (Vector Laboratories) diluted 1:1000, DAPI (Sigma) diluted 1:1000, and secondary antibodies diluted 1:1000 in PBS + 3% BSA + 0.1% TritonX-100 filtered through a 0.2um filter for either 1.5-2 hours at room temperature or overnight at 4C: Alexa Fluor 568 donkey α rabbit (Molecular Probes), Alexa Fluor 647 donkey α rabbit (Molecular Probes). Finally, the secondary antibodies were washed out with PBS three times for 10 minutes each, covered in a drop of VectaShield (Vector Laboratories) and a 25×50mm coverslip (Thermo Fisher), and sealed with nail polish.

### Immunofluorescence on whole mount testes

Testicular tubules from adult *Cenpa*^*mScarlet/+*^ male mice were prepared as previously published with some modifications (Smith and Braun, 2012). Briefly, the tunica albuginea was removed from the testes of euthanized males and the seminiferous tubules were teased apart in PBS. Interstitial cells were removed by incubating the tubules in 0.5mg/mL collagenase (Sigma) at room temperature (RT) for 5 minutes. The tubules were then rinsed three times with PBS, and fixed in 2% paraformaldehyde (Electron Microscopy Sciences) for 6 hours at 4°C. Paraformaldehyde-fixed tubules were then rinsed three times with PBS, and permeabilized with 0.25% NP-40 (US Biologicals) in PBST (PBS + 0.05% Tween (Thermo Fisher) for 25 minutes at RT. The tubules were rinsed three times for 5 minutes each with PBST and blocked using 5% normal donkey serum (Jackson Immuno Research) in PBST either 2 hours at RT or over-night at 4°C. The tubules were incubated with one of the following primary antibodies diluted in PBST + 5% normal donkey serum at 4C overnight: rabbit α CENP-A at 1:250 (Cell Signalling Technologies 2048), rabbit α CENP-A-Ser18ph at 1:100 (Active Motif 61483), Gt GFRa at 1:250 (R&D Systems AF429), Rb Sohlh1 at 1:200 (gift from Aleksandar Rajkovic), Rb Stra8 at 1:250 (Abcam ab49602), or mouse gH2A.X at 1:500 (EMD Millipore 05-636). The next day, the tubules were washed three times for 10 minutes each in PBST, then stained with DAPI at a 1:1000 dilution and an experiment-dependant combination of the following secondary antibodies at room temperature for 1 hour in PBST + 5% normal donkey serum: Alexa Fluor 488 donkey α goat at 1:500 (Molecular Probes), Alexa Fluor 488 donkey α mouse at 1:1000 (Molecular Probes), Alexa Fluor 568 donkey α rabbit at 1:1000 (Molecular Probes), Alexa Fluor 568 donkey α mouse at 1:1000 (Molecular Probes), Alexa Fluor 647 donkey α rabbit at 1:1000 (Molecular Probes), and/or Alexa Fluor 647 donkey α mouse at 1:1000 (Molecular Probes). After secondary antibody incubation, the tubules were rinsed three times for 10 minutes each with PBST. The tubules were then gently spread onto a coverslip with a paintbrush, dried briefly, covered in a drop of VectaShield (Vector Laboratories) and a 25×50mm coverslip (Thermo Fisher), and sealed with nail polish.

### Preparation of spermatocyte spreads

Spermatocytes were mounted and spread onto glass slides as previously published with some modifications (Zelazowski et al., 2017). Briefly, a single testis from an adult C57Bl/6 or *Cenpa*^*mScarlet/+*^ male was decapsulated by shaking at 150rpm for 55 minutes at 32C in testis isolation medium (TIM) comprised of 104 mM NaCl (Sigma) + 45 mM KCl (Sigma) + 1.2 mM MgSO4 (Sigma) + 0.6 mM KH2PO4 (Sigma) + 0.1% (w/v) glucose (Sigma) + 6 mM sodium lactate (Sigma) + 1 mM sodium pyruvate pH 7.3 (Sigma) with an additional 2mg/mL collagenase (Worthington) added for this step only. The tubules were washed three times in TIM at room temperature before being resuspended in a small volume of TIM + 0.7 mg/mL Trypsin (Sigma) + 4 mg/mL DNaseI (Roche 104159) and shaken at 150rpm for 15 minutes at 32C. The trypsin digest was stopped by diluting Trypsin Inhibitor (Sigma T9003) to 5 mg/mL and DNaseI to 0.1 ug/uL in the resulting single cell suspension and cell clumps were separated by gentle pipetting with a plastic transfer pipet and filtering through at 70um mesh filter (Thermo Fisher). The cells were then washed once each in TIM + 1:1000 DNaseI and PBS + 1:1000 DNaseI with centrifugation at 500g. The dissociated cells were then separated into twelve equal aliquots and two aliquots were spread onto glass slides in turns. Each aliquot was resuspended in pre-warmed 80uL 0.1 M sucrose (Sigma) and incubated for 3-5 minutes at 37C, then 20uL of the aliquot was slowly spread along the length of a positively charged and precleaned glass slide. Each aliquot was spread onto four glass slides, which were each pre-covered with 65uL of prewarmed 1% paraformaldehyde pH 9.2 (EMD Millipore) + 0.1% TritonX-100 during the aliquot’s 3-5 minute incubation in 0.1 M sucrose. After spreading all of the aliquots of the testicular single cell suspension onto glass slides, the slides were allowed to dry slowly at room temperature for 3.5 hours. Finally, the slides were washed twice for five minutes each in PBS + 0.004% Photo-Flo 200 (Kodak 1464510), air dried, and stored at −80C.

### Immunofluorescence staining of spermatocyte spreads

Spermatocytes spreads were immunostained following a previously published protocol with some modifications (Zelazowski et al., 2017). Briefly, the desired number of slides were removed from storage at −80C and dried for 10-20 minutes at room temperature, then immediately blocked for 30 minutes at room temperature in PBS (Thermo Fisher) + 1% fetal bovine serum (FBS) (Thermo Fisher) + 0.3% bovine serum albumin (BSA) (Sigma) + 0.05% Triton X-100 (Sigma) and incubated with primary antibodies diluted in PBS + 1% FBS + 0.3% BSA + 0.05% Triton X-100 overnight at room temperature: rabbit α CENP-A at 1:250 (Cell Signalling Technologies 2048), rabbit α Sycp1 at 1:200 (Abcam ab15090), or mouse α gH2A.X at 1:5000 (EMD Millipore 05-636). The next day, the spreads were washed with PBS + 1% FBS + 0.3% BSA + 0.05% Triton X-100 four times for 1 minute, 5 minutes, 10 minutes, and 15 minutes respectively, all at room temperature. Next, the spreads were stained with DAPI diluted 1:500 and secondary antibodies diluted 1:200 in PBS + 1% FBS + 0.3% BSA + 0.05% Triton X-100 for 45 minutes at 37C in a humidified chamber: Alexa Fluor 488 donkey α mouse (Molecular Probes), and Alexa Fluor 647 donkey α rabbit (Molecular Probes). The spreads were then washed again with PBS + 1% FBS + 0.3% BSA + 0.05% Triton X-100 four times for 1 minute, 5 minutes, 10 minutes, and 15 minutes respectively, all at room temperature. Finally, they were washed twice for five minutes each in PBS + 0.004% Photo-Flo 200 (Kodak 1464510), dried briefly, covered in a drop of VectaShield (Vector Laboratories) and a 25×50mm coverslip (Thermo Fisher), and sealed with nail polish.

### Flow cytometry

Testes from adult *Cenpa*^*mScarlet/+*^ male mice were dissociated, stained, and flow sorted as previously published with some modifications (Green et al., 2018). Briefly, the tunica albuginea was removed from the testes of euthanized males and the seminiferous tubules were transferred to 10ml of Advanced DMEM:F12 media (Thermo Fisher) + 200 mg/ml Collagenase IA (Sigma) + 400 units/mL DNase I (Worthington Biochemical Corp). Tubules were dispersed by gently shaking and allowed to settle for 1 minute at room temperature. Excess media was removed with a sterile pipette, leaving ∼2mL with the settled tubules. The tubules were then dissociated at 35C with horizontal shaking at 215 rpm for 10 minutes in an additional 10mL of Advanced DMEM:F12 media (Thermo Fisher) + 400 units/mL DNase I (Worthington Biochemical Corp) + 200 mg/ml Trypsin (Thermo Fisher). The trypsin was quenched with the addition of 3mL of fetal bovine serum (FBS) (Thermo Fisher). The resulting single cell suspension was filtered through a 100um strainer (Thermo Fisher), washed in PBS (Thermo Fisher), pelleted at 600g for 3 minutes, and re-suspended in 6mL MACS buffer containing 0.5% BSA (Miltenyi Biotec). The suspension was stained with Hoechst 3342 (Thermo Fisher) and propidium iodide (PI) (Thermo Fisher) as previously published with no modifications (Gaysinskaya et al., 2014). Spermatogonia (2n), spermatocytes (4n, mostly pachytene and diplotene), and round spermatids (1n) were isolated from the stained live single cell suspension based on DNA content and DNA compaction using a FACS ARIA II/III flow cytometer (BD Biosciences) available at the University of Michigan’s Flow Cytometry Core. Gates and sorting conditions were adapted from previously published methods and optimized for each biological replicate (Gaysinskaya et al., 2014; Green et al., 2018). Sorted cell populations were immediately pelleted at 600g and snap frozen in liquid nitrogen before storage at −80C.

### Sperm chromatin immunoprecipitations

Mouse sperm was collected in a swim up from the caudal epididymis and vas deferens as previously described (Brykczynska et al., 2010), and mature human sperm was collected and snap frozen as detailed above. Chromatin from both human and mouse sperm was immunoprecipitated using a previously published protocol with some modifications (Yoshida et al., 2018). Briefly, 30 million mature sperm were resuspended in 1mL of PBS and cross-linked by 1% formaldehyde (Sigma) for 10□minutes at room temperature before quenching the fixation by adding Tris pH 7.5 to a final concentration of 0.2 M and incubating for another 10 minutes at room temperature. The fixed sperm was then washed twice in sperm decondensation buffer: 5□mM HEPES pH 8.0 (Sigma) + 1□mM PMSF (Sigma) + 0.2% NP-40 (Sigma) + 10□mM EDTA (Sigma) + 5□mM NaCl (Sigma) + 1.2 M urea (Sigma) + 10□mM DTT (Sigma) + 2XLcomplete protease inhibitor cocktail (Sigma). After washing, the sperm was resuspended in sperm decondensation buffer with 1 mg/mL heparin sodium salt (Sigma) at a concentration of 15 million sperm per 3 mL decondensation buffer. The mouse sperm was then decondensed by incubating for 5 hours at 42C, while human sperm was decondensed by incubating for 2 hours at 42C. The decondensed sperm was then washed twice in lysis buffer: 50□mM HEPES pH 7.5 (Sigma) + 140□mM NaCl (Sigma) + 1□mM EDTA (Sigma) + 10% glycerol (Sigma) + 0.5% NP-40 (Sigma) + 0.25% Triton X-100 (Sigma) + and 1X□complete protease inhibitor cocktail (Sigma). After washing, decondensed mouse sperm was treated with lambda phosphatase (NEB) by resuspending the sperm in 1X NEBuffer for Protein MetalloPhosphatases supplemented with 1 mM MnCl2 (NEB) and incubating for 30 minutes at 37C. The decondensed human sperm was not treated with lambda phosphatase. Both mouse and human sperm were then resuspended in 50□mM Tris-HCl pH 8.0 (Sigma) + 10□mM EDTA (Sigma) + 1% SDS (Sigma) + 1X complete protease inhibitors (Sigma) + 1□mM PMSF and sonicated using four cycles of 30 seconds on/off. The sonicated sperm was diluted 1:10 in 10□mM Tris-HCl pH 8.0 (Sigma) + 100□mM NaCl (Sigma) + 1□mM EDTA (Sigma) + 0.5□mM EGTA (Sigma) + 1% Triton X-100 (Sigma) + 0.1% sodium deoxycholate (Sigma) + and 1XLcomplete protease inhibitor cocktail (Sigma) to dilute SDS to a final concentration 0.1%. Next, the sperm chromatin was spun down for 10 minutes at 20,000g and the insoluble fraction was saved for immunoblot analysis. The soluble chromatin was precleared by gently rotating with magnetic Protein A beads (Invitrogen 10002D) for 1 hour at 4C, pre-clearing beads were removed and 10% of the soluble chromatin was saved as input for immunoblot analysis. Chromatin was immunoprecipitated by incubating the soluble chromatin with the desired antibody pre-bound to magnetic Protein A beads for at least 14 hours at 4C with gentle rotation. Antibodies used for immunoprecipitation include rabbit α CENP-A at 1:200 (Cell Signalling Technologies 2048) for mouse sperm, rabbit α HJURP at 1 ug per 10 million sperm (Abcam ab100800) for human sperm, and rabbit IgG isotype control at either 1:200 or 1 ug per 10million sperm (Invitrogen 10500C) for mouse or human sperm respectively. The unbound fraction was saved for immunoblot analysis, and the immunoprecipitation was washed four times in 50 mM HEPES pH 7.0 (Sigma) + 0.5 M LiCl (Sigma) + 1 mM EDTA (Sigma) + 0.7% sodium deoxycholate (Sigma) + 1% NP-40 (Sigma), then twice in 10 mM Tris-HCl pH 8.0 (Sigma) + 1 mM EDTA (Sigma), and the immunoprecipitation was finally eluted in 10 mM Tris-HCl pH 8.0 (Sigma) + 1 mM EDTA (Sigma) + 1% SDS (Sigma) + 1X□complete protease inhibitor cocktail (Sigma). The input (10% of IP), 10% of the unbound, and 100% of the elution were immunoblotted from each experiment and antibody. Immunoblotted membranes were incubated with either rabbit α HJURP at 1:500 (Abcam ab100800), rabbit α CENP-A at 1:250 (Cell Signalling Technologies 2048), mouse α CENP-A at 1:1000 (GeneTex GTX13939), mouse α H2B at 1:500 (Abcam ab52484), or rabbit α H2B at 1:1000 (Cell Signalling Technologies 12364). Primary antibodies hosted in mouse were detected with HRP-conjugated goat α mouse IgG at 1:10,000 (Abcam ab6721) and primary antibodies hosted in rabbit were detected using HRP conjugated mouse α rabbit IgG light chain at 1:5,000 (Abcam ab99697).

### In-vitro fertilization and drug treatments

Mouse zygotes and early embryos were collected by in vitro fertilization (IVF) with oocytes collected from either CF-1 female mice (Charles River Laboratories), C57Bl/6J females (Jax), or *Cenpa*^*mScarlet/+*^ females and sperm collected from either B6D2 males (Jax), or *Cenpa*^*mScarlet/+*^ males. *Cenpa*^*mScarlet/+*^ males were allowed to mate once before the IVF and both B6D2 and *Cenpa*^*mScarlet/+*^ males were housed individually for seven days prior to IVF. Males were 8-12 weeks of age and females 3-4 or 6-10 weeks of age at the time of gamete collection. Females were superovulated with an intraperitoneal injection of 100 uL HyperOva (Card KYD-010-EX-X5) and 5-7.5 IU of human chorionic gonadotropin (hCG) (Sigma CG5) at 60 and 14 hours respectively before oocyte collection. Media for sperm capacitation and IVF incubation, and if applicable later stage embryo maturation, was allowed to equilibrate in an incubator at 37C with 5% CO2 the night before IVF. The day of IVF, males were euthanized by cervical dislocation 1.25 hours before oocyte collection. Mature sperm was collected by mincing the epididymis and vas deferens and capacitating in 500 uL Research Vitro Fert Media (Cook Medical K-RVF-50) for 10 minutes at 37C and 5% CO2. The epididymis and vas deferens tissue were then removed, sperm concentration and percent motility were counted using a Makler sperm counter (Cooper Surgical SEF-MAKL), and sperm was allowed to capacitate for another 1hr. Females were euthanized by cervical dislocation exactly 14 hours post hCG injection and cumulus-oocyte complexes were extracted from the oviduct in pre-warmed MEM media (Thermo Fisher 12360038) and immediately transferred into 500 uL Research Vitro Fert Media and incubated with 1 million motile sperm. Fertilization was allowed to continue for 4-6 hours before the zygotes were removed from the well containing sperm and placed in either another pre-equilibrated 500 uL of Research Vitro Fert Media for collection at the 1-cell stage or washed 5 times and incubated in pre-equilibrated KSOM AA (Millipore Sigma) at 37C and 5% CO2 and allowed to develop to morula or blastocyst stage embryos. Transcription-inhibited zygotes were treated with 5,6-dichloro-1-β-D ribofuranosyl-benzimidazole (Sigma) diluted 1:200 to 200 uM and 5-ethynyl uridine (Sigma) diluted 1:500 to 500 nM, both added to the IVF media 1.5 hours after fertilization, embryos were collected 10 hours after fertilization. Translation-inhibited zygotes were treated with cycloheximide (Sigma) diluted 1:500 to 100 ug/mL starting 1 hour after fertilization and O-propargyl-puromycin (Click Chemistry Tools 1493) diluted 1:500 to 20 uM in the IVF media 1 hour before collecting the zygotes 8.5 hours after fertilization. Zygotes with DNA replication inhibited were treated with aphidicolin (Sigma) diluted 1:500 to 10 ug/mL starting 1.5 hours after fertilization and 5-Ethynyl 2’-deoxyuridine (Sigma) diluted 1:500 to 20 uM added 1 hour before collection at 8.5 hours after fertilization. Zygotes cultured for controls were collected and fertilized alongside every replicate of the drug treatment experiments, and were stained with either EU, OPP, or EdU in the same manner. However, the control zygotes had dimethyl sulfoxide (Sigma) added to their IVF media at the same dilutions and time course as those zygotes treated with a small molecule inhibitor.

### Immunofluorescence on oocytes and early embryos

For immunofluorescence, embryos or oocytes were collected at the indicated timepoints after washing in M2 media (Thermo Fisher), treated briefly (30 seconds-1 minute) with Acidic Tyrodes solution (EMD Millipore) to remove the zona pellucida, and fixed in 4% paraformaldehyde (EMD Millipore) + PBS (Thermo Fisher) + 0.04% TritonX-100 (Sigma) + 0.3% Tween-20 (Sigma) + 0.2% sucrose (Sigma) for 10 minutes at 37C. Fixed embryos or oocytes were stored under mineral oil (Sigma) in PBS at 4C for no longer than one week. All staining took place in nine-well depression Pyrex plates (Millipore Sigma CLS722085) and utilized buffers which were made fresh and filtered through 0.2 um PES filters (Whatman 6780-2502) each day of staining. On the first day of immunofluorescence staining the embryos or oocytes were permeabilized in PBS + 0.5% TritonX-100 (Sigma) + 3% bovine serum albumin (BSA) (Sigma) at room temperature for 1 hour. If the embryos were incubated with EdU, 5-EU, or OPP (see above), these reagents, which each contain a terminal alkyne group, were next conjugated to a AZDye 488 picolyl azide following the Click Chemistry Tools kit protocol (CCT #1493). If no additional immunostaining was necessary the treated embryos were stained with DAPI (Sigma) diluted at 1:200 in PBS + 0.1% Triton + 3% BSA for 15-30 minutes at room temperature, washed twice for 10 minutes each with PBS + 0.1% Triton + 3% BSA, briefly dried onto a glass slide, then covered in Vectashield (Vector Laboratories) and a 22×22mm no. 1.5H coverslip (Thermo Fisher). After permeabilization and optional click-chemistry staining, embryos or oocytes were blocked in PBS + 0.1% Triton + 3% BSA + 10% fetal bovine serum (Thermo Fisher) at room temperature for 1-2 hours and stained with primary antibodies diluted in blocking buffer at 4C overnight: rabbit α CENP-A at 1:250 dilution (Cell Signalling Technologies 2048), or rabbit α CENP-C at 1:1000 dilution (gift from Ben Black). Embryos or oocytes were washed the following day in PBS with 0.1% Triton and 3% BSA five times for at least 15 minutes each, followed by incubation with DAPI at 1:200 and secondary antibodies diluted 1:500 in PBS + 0.1% Triton + 3% BSA + 10% fetal bovine serum for 2 hours at room temperature: Alexa Fluor 488 donkey α rabbit (Molecular Probes), Alexa Fluor 568 donkey α rabbit (Molecular Probes). The stained embryos or oocytes were washed thrice for 10 minutes each with PBS + 0.1% Triton + 3% BSA, briefly dried onto a glass slide, then covered in Vectashield and a 22×22 mm no. 1.5H coverslip.

### Oocyte collection and maturation

Germinal vesicle (GV) stage mouse oocytes were collected from either CF-1 females (Charles River Laboratories), C57Bl/6J females (Jax), or *Cenpa*^*mScarlet/+*^ females at 3-4 or 6-10 weeks of age. The females were each injected with 5-7.5 IU pregnant mare’s serum gonadotropin (Sigma) intraperitoneally 44-48 hours prior to sacrifice. Media for oocyte collection and maturation were placed in an incubator to equilibrate to 37C and 5% CO2 the day before oocyte collection. Milrinone (Sigma) and fetal bovine serum (Thermo Fisher) were diluted to 2.5 uM and 5% respectively in both oocyte and collection media an hour before oocyte collection. Females were euthanized by cervical dislocation and ovaries were dissected out and placed in MEM (Thermo Fisher 12360038) + 2.5 uM milrinone + 5% FBS. Oocyte-cumulus cell complexes were isolated by repeatedly poking the ovaries’ antral follicles with an insulin syringe. Oocytes were washed through several drops of MEM and cumulus cells were mechanically detached by repeated mouth pipetting the complexes up and down. Cleaned GV oocytes were then washed and allowed to recover in MEMa with GlutaMAX (Thermo Fisher 32561-037) + 2.5 uM milrinone + 5% FBS for several hours in 37C and 5% CO2. The GV oocytes were washed away from milrinone by transferring them through at least 5 drops of MEMa with GlutaMAX (Thermo Fisher 32561-037) + 5% FBS, at which point they were then cultured at 37C and 5% CO2. Metaphase I (MI) oocytes were collected 8 hours after release from milrinone and metaphase II (MII) oocytes were collected 16 hours after release from milrinone. If the MII eggs were to be fertilized, they were transferred at least five times through drops of pre-equilibrated Research Vitro Fert Media before being fertilized in 500 uL Research Vitro Fert Media with 2 million motile sperm. All zygotes from in-vitro matured and in-vitro fertilized oocytes were collected after 8.5 hours of fertilization.

### Oocyte microinjection

GV oocytes were collected as described above and microinjected according to previously published protocols (Stein and Schindler, 2011). Briefly, after letting GV oocytes recover from collection, the oocytes were transferred into MEM (Thermo Fisher 12360038) + 2.5 uM milrinone (Sigma) + 5% FBS (Thermo Fisher) + 1% bovine serum albumin (Sigma) and injected in groups of 10 with a FemtoJet 4i microinjector (Eppendorf). Holding micropipettes (Cooper Surgical MPH-MED-35) were cleaned with 70% ethanol (Sigma) and ddH2O, and injecting needles (Sutter B100-75-10) were pulled with a Sutter P97 micropipette fuller fresh before every injection. RNA was prepared for injection by first linearizing the pIVT plasmids with PvuI (NEB) and pGEMHE plasmids with AscI (NEB), 100-500ng was then utilized as the template for a T7 in vitro transcription reaction with the mMessage mMachine T7 kit (Thermo Fisher AM1344). Finally, the RNA was purified with AMPure XP beads at a 2:1 bead:RNA ratio, aliquoted and stored at −80C. While the collected GV oocytes were recovering, RNA integrity was checked by denaturing gel electrophoresis and good quality IVT RNA was combined for the desired injection combination, diluted to 100 ng/uL in dH2O, and spun down at 20,000g for 30min at 4C. Injected GV oocytes were allowed to recover in MEMa with GlutaMAX (Thermo Fisher 32561-037) + 2.5 uM milrinone + 5% FBS for 2-6 hours at 37C and 5% CO2. The injected oocytes were matured and fertilized as described above. All zygotes from injected, in-vitro matured, and in-vitro fertilized oocytes were collected after 8.5 hours of fertilization.

### Oocyte photobleaching

GV oocytes were collected as described above and imaged and photobleached according to previously published protocols (Christodoulou et al., 2019; Ooga and Wakayama, 2017). Briefly, after recovering from collection, the GV oocytes were stained with 10ug/mL Hoechst 3324 (Sigma) for 30 minutes and then individually transferred into an opening in a 105um nylon mesh net (Tisch Scientific ME17250) affixed to the bottom of a well in an 8-well glass bottom µ-Slide (Ibidi 80807) with silicone (Rutland). They were incubated in MEMa with GlutaMAX (Thermo Fisher 32561-037) + 2.5 uM milrinone (Sigma) + 5% FBS (Thermo Fisher) pre-equilibrated to 37C at 5% CO2. The GV oocytes were incubated in a OkoLab live-cell imaging chamber (H301) on a Nikon Nikon A1R with a 60X 1.2NA long range water-immersion objective available at the University of Michigan’s Center for Live Cell Imaging through the Microscopy and Image Analysis Laboratory. Nikon NIS Elements was used to photobleach the mScarlet fluorescence in the nucleus of each GV oocyte with a circular ROI, the area adjusted to the size of each nucleus, with a 561nm laser line at 90uW of power and a 19.9uSec pixel dwell time. After photobleaching, the oocytes were allowed to recover for at least 2 hours before being in-vitro matured and in-vitro fertilized as described above. All zygotes from photobleached, in-vitro matured, and in-vitro fertilized oocytes were collected after 8.5 hours of fertilization.

## QUANTIFICATION AND STATISTICAL ANALYSIS

### Quantification of centromere signals

Quantification of centromere fluorescence intensity was made in Fiji/ImageJ software by drawing circles of constant diameter around individual centromeres across all Z-planes of the cell. The total intensity was calculated for each centromere puncta after subtracting the background of the image, which was obtained from areas of corner regions of each image, and then summing the total puncta fluorescence intensities for each pronuclei or cell being studied. For each statistical test, multiple independent technical and biological replicates were quantified to obtain a minimum of ten embryos per experiment.

### Statistical Testing

All statistical tests for significance were performed in either R or GraphPad Prism 9 software. P-values were calculated at a significance level of 0.05 or 95% confidence interval for all two tailed statistical tests. Figures were made in R and GraphPad Prism 9 software. The number of replicates for each experiment is supplied in the figure legends. Tests for normal distribution of the data were performed in R using both the shapiro.test() and leveneTest() functions. Normally distributed paired data was tested using the t.test() and non-parametric data was tested using the wilcox.test(). In GraphPad, the Shapiro-Wilk normality test was applied to determine if the data was normally distributed, and the appropriate normal or non-parametric t-test was applied to the data accordingly.

**Supplemental Figure 1: Generation and validation of *Cenpa***^***mScarlet/+***^ **mice**

A&B) Testes-to-body weight ratio (A) and total sperm count (B) are maintained in *Cenpa*^*mScarlet/+*^ mice. Each dot is the measurement from one mouse. ns indicates not significant. Mean values for testes/body weight ratio are as follows: WT = 0.3732 from n = 4 mice, *Cenpa*^*mScarlet/+*^ = 0.3644 from n = 6 mice. Mean values for sperm count are as follows: WT = 28.25million from n = 4 mice, *Cenpa*^*mScarlet/+*^ = 29.5million from n = 6 mice. C) The colocalization of CREST and direct CENP-A-mScarlet fluorescence in *Cenpa*^*mScarlet/+*^ testes further supports that the CENP-A tag does not affect localization. Representative images from n = 3 staining experiments and n = 2 mice. Scale bar is 20um. D) Immunostaining of *Cenpa*^*mScarlet/+*^ spermatocyte spreads with SYCP3 confirms mScarlet enrichment at chromosome ends, consistent with murine telocentric chromosomes and lack of ectopic CENP-A deposition. Scale bar is 10um. E) Quantification of western band intensities from Fig 1C, normalized to ploidy and graphed as the percentage of spermatogonial protein levels. Mean values for H3 are as follows: spermatogonia = 100%, spermatocyte = 96.46%, spermatid = 41.77%, sperm = 1.26%. Mean values for V5 are as follows: spermatogonia = 100%, spermatocytes = 54.50%, spermatids = 6.50%, sperm = 4.59%. Mean values for total CENP-A are as follows: spermatogonia = 100%, spermatocytes = 61.77%, spermatids = 18.28%, sperm = 20.38%. Mean values for CENP-B are as follows: spermatogonia = 100%, spermatocytes = 24.94%, spermatids = 36.46%, sperm = 1.76%. Mean values for CENP-C are as follows: spermatogonia = 100%, spermatocytes = 307.02%, spermatids = 20.15%, sperm = 0%. Mean values for HJURP are as follows: spermatogonia = 100%, spermatocytes = 106.83%, spermatids = 28.35%, sperm = 3.88%. F) Immunoblots of CENP-B protein levels in flow sorted germ cells and mature sperm from C57Bl/6J testes. Representative image from n = 2 replicates from n = 2 mice.

**Supplemental Figure 2: The increase in CENP-A density is a conserved property in *Drosophila* GSCs**

A) Quantification of endogenous ID4-EGFP fluorescence in all (n = 157) A_s_ or A_pr_ EGFP+ cells. ID4-EGFP high and low cutoffs are marked as horizontal dotted lines and were defined as the cells in the top or bottom 33 percentile of fluorescence, the top cutoff is ≥992.4au and the bottom is ≤601.51au. B) Quantification of endogenous ID4-EGFP fluorescence in A_single_ (As) and A_paired_ (Ap) cells. n_As_ = 71, n_Ap_ = 86. Mean fluorescence of ID4-EGFP in As and Ap are: As = 1027.11, Ap = 773.31. C) Quantification of total CENP-A immunofluorescence in ID4-EGFP high and ID4-EGFP low cells in As and Ap populations. n_Id4 high_ = 52, n_Id4 low_ = 52. Mean fluorescence of CENP-A in ID4-EGFP high and ID4-EGFP low is as follows: CENP-A_Id4 high_ = 912.94, CENP-A_Id4 low_ = 644.36. D) A schematic diagram of *Drosophila melanogaster* spermatogenesis. E) Representative images of each step of spermatogenesis. Germ cells expressing Tubulin-GFP and immunostained with anti-CENP-A^CID^ antibody. Asterisk: hub. Scale bars are 5um. F) Quantification of total fluorescence of CENP-A^CID^ across the stages of spermatogenesis. Mean fluorescence values are as follows: hub cells = 154084.1±10361.88 (n=20), Germline stem cells (GSCs) = 540812.79 ±32852.42 (n=34), spermatogonia cells (SGs) 4-cells = 435385.08 ±34884.65 (n=13), SGs 8-cells = 343275.60 ±20491.73 (n=33), SGs 16-cells = 337345.31 ±18628.79 (n=26), and primary spermatocytes (SCs) = 313588.84 ±13514.78 (n=19); *\**: p<0.05, *\*\**; p<0.01, *\*\*\*\**: p<0.0001. All values = mean ± SE.

**Supplemental Figure 3: CENP-A levels increase during meiotic prophase I in oocytes and are not actively restored during the MI-MII transition**

A) Representative images of total CENP-A immunofluorescence in OCT4-EGFP fetal gonads collected at the indicated embryonic and adult timepoints (eg. E11.5 is 11.5 days post conception). Scale bars are 20um, oogonia/oocytes are circled in yellow. Images are representative of n = 3 technical replicates from two gonads collected at each embryonic timepoint from one pregnant female mouse. B&C) Quantification of total CENP-A (B) and CENP-C (C) immunofluorescence at the indicated embryonic and adult timepoint. Each dot is the sum of CENP-A-mScarlet or CENP-C puncta in one cell or pronucleus. Mean intensity of the total CENP-A fluorescence is as follows: Somatic = 397au, n = 41 cells, E11.5 = 3298au from n = 32 PGCs, E13.5 = 2383au from n = 51 oogonia/oocytes, E18.5 = 3606au from n = 50 oocytes, P2 = 2688au from n = 30 oocytes, GV = 10866au from n = 13 oocytes. Mean intensity of the total CENP-C fluorescence is as follows: Somatic = 2234au from n = 26 cells, E11.5 = 6505au from n = 38 PGCs, E13.5 = 3770au from n = 31 oogonia/oocytes, E18.5 = 5777au from n = 42 oocytes, P2 = 3208au from n = 50 oocytes, GV = 13753au from n = 14 oocytes. D) Representative images of endogenous CENP-A-mScarlet fluorescence in geminal vesicle (GV), metaphase I (MI) and metaphase II (MII) stage eggs. Oocytes were collected from n = 6 *Cenpa*^*mScarlet/+*^ females, and *in vitro* matured in n = 3 replicates. Scale bar is 10um. E&F) Quantification of endogenous CENP-A-mScarlet fluorescence (E) or CENP-C immunofluorescence (F) at each of the oocyte stages. Each dot is the sum of CENP-A-mScarlet or CENP-C puncta in one cell or pronucleus. Mean intensity of the total CENP-A fluorescence is as follows: GV = 1544au from n = 55 oocytes, MI = 1814au from n = 28 oocytes, MII polar body = 913.5au from n = 21 polar bodies, MII oocyte = 1068au from n = 21 oocytes. Mean intensity of the CENP-C immunofluorescence is as follows: GV = 6556au from n = 42 oocytes, MI = 10608au from n = 28 oocytes, MII polar body = 8794au from n = 10 polar bodies, MII oocyte = 14347au from n = 10 oocytes, PN0 polar body = 5192au from n = 10 polar bodies. *: p < 0.05, **: p < 0.01, ***: p < 0.001, ****: p < 0.0001. ns indicates not significant.

**Supplemental Figure 4: Centromere epigenetic memory is cell-type and sex specific**

A) Immunoblots of CENP-A and H3 protein levels in mouse TG2a embryonic stem cells (ESCs) to mature mouse sperm. Shown is a representative immunoblot membrane from n = 3 immunoblots on n = 3 mice. B) Immunoblots of CENP-A and H3 protein levels from human embryonic stem cells (ESCs) and human mature sperm. Shown is a representative immunoblot from n = 2 immunoblots on n = 2 deidentified human sperm samples. C) Relative quantification of the abundance of CENP-A or H3 protein retained in Human and Mouse sperm. Graphs are the average percent of CENP-A or H3 band intensity from sperm vs. ESCs, normalized to cell input. Human H3 mean = 3.125%, human CENP-A mean = 12.5%, mouse H3 mean = 1.5625%, mouse CENP-A mean = 18.75%. D) Immunoblot of protein from GV and MII stage oocytes alongside mature sperm with the indicated cells loaded per lane. The membrane was stripped and re-probed with the indicated antibodies. Representative image from n = 2 blots using oocytes from n = 12 females and n = 2 males. E) Quantification of western band intensities from D). Data is shown normalized to ploidy, cellular input, and GV protein levels. Mean values are as follows: GV CENP-A = 100, MII CENP-A = 165.26, Sperm CENP-A = 17.5, GV CENP-C = 100, MII CENP-C = 183.7, Sperm CENP-C = 0. F) Quantification of Cenpa RNA transcripts from *in vitro* matured GV and MII eggs, early (6hpf) and late (12hpf) zygotes, and 2-, 4-, and 8-cell embryos. RNA levels are shown as a percentage of GV: TG2a = 13.42%, GV = 100%, MII = 53.23%, Early Zygote = 32.13%, Late Zygote = 45.83%, 2-cell = 132.5%, 4-cell = 256.92%, 8-cell = 20.43%. n = 2 RT-qPCRs were run on oocytes and embryos from n = 6 mice.

**Supplemental Figure 5: Progression through the early pronuclear stages is not affected by drug treatments**

A-C) Quantification of CENP-A-mScarlet direct fluorescence in each experimental replicate for each of the drug treatments shown in Fig 5. Each color represents data from one experimental replicate. Each dot is the sum of CENP-A-mScarlet puncta in one pronucleus. D-F) Quantification of pronuclear stages in zygotes treated with either DRB (D), CHX (E), Aph (F). G) Representative images from 2-cell embryos treated with DRB and EU as in Figure 5A show full inhibition of zygotic genome activation. Images are representative of n = 2 IVFs using n = 6 total females. Scale bar is 20um. H) Quantification of CENP-A-mScarlet direct fluorescence from only PN3 stage embryos treated with cycloheximide. Each dot is the sum of CENP-A-mScarlet puncta in one pronucleus. *: p < 0.05, ns indicates not significant. Mean fluorescence intensities shown in H) are as follows: Control maternal = 3515au from n = 12 embryos, Control paternal = 926.5au from n = 12 embryos, CHX+ maternal = 2496au from n = 37 embryos, CHX+ paternal = 585au from n=33 embryos.

**Supplemental Figure 6: CENP-A levels in zygotes are predetermined by the available pool of CENP-A and deposition machinery**

A) Representative images of CENP-A-Emerald and total CENP-A in GV injected (CF1) oocytes that are *in vitro* maturated and/or *in vitro* fertilized. Oocytes and embryos were immunostained for total CENP-A and imaged along with the direct fluorescence from CENP-A-Emerald. Scale bars are 20um in the main figure and 5um in the PN0 and PN3 insets. B) Quantification of total CENP-A immunofluorescence intensity from control, CENP-A-Emerald-only injected, or CENP-A-Emerald + mCherry-HJURP injected oocytes. Note that only embryos at PN3-5 were included in the quantification. Each dot is the sum of total CENP-A puncta in one pronucleus. *: p < 0.05, ****: p < 0.0001. ns indicates not significant. Mean fluorescence intensities shown in B) are as follows: Control maternal = 6981au from n=54 embryos, Control paternal = 6922au from n = 54 embryos, injected CENP-A-Emerald maternal = 18727au from n = 33 embryos, injected CENP-A-Emerald paternal = 22495au from n = 33 embryos, injected CENP-A-Emerald + mCherry-HJURP maternal = 21984au from n = 26 embryos, injected CENP-A-Emerald + mCherry-HJURP paternal = 23819au from n = 26 embryos. Images and quantification come from n = 7 injection experiments using n = 12 CF1 females.

**Supplemental Figure 7: Asymmetric deposition of CENP-A is correlated with asymmetric levels of CENP-C and MIS18BP1**

A) Immunofluorescence images of CENP-C and direct fluorescence from maternal CENP-A-mScarlet at the indicated zygotic pronuclear stages (PN). Images are representative of n = 2 IVFs using n = 6 *Cenpa*^*mScarlet/+*^ females and n = 1 (C57Bl/6J X DBA2) F1 males. All scale bars are 20um. B) Quantification of CENP-C immunofluorescence shown in panel A, grouped by pronuclei and pronuclear stage. Each dot is the sum of CENP-C puncta in one pronucleus. **: p < 0.01. Mean fluorescence intensities are as follows: Maternal PN0-2 = 10658 from n = 34 pronuclei, Paternal PN0-2 = 15661 from n = 34 pronuclei, Maternal PN3-5 = 6673 from n = 89 pronuclei, Paternal PN3-5 = 10014 from n = 89 pronuclei. C) Immunofluorescence images of MIS18BP1 across the pronuclear stages showing MIS18BP1 is localized to pronuclei beginning at PN2 and is enriched in paternal chromatin. Representative images from n = 3 IVFs using n = 7 females. The scale bars are 20um. D) PN4 zygotes immunostained for MIS18BP1 imaged along with maternal CENP-A-mScarlet fluorescence. Pronuclear insets reveal that MIS18BP1 co-localizes with CENP-A-mScarlet, and this colocalization only occurs in the paternal pronucleus (arrowheads). Representative images from n = 1 IVF using n = 3 females. The scale bars are 20um. PN = pronuclear stage. E) Quantification of pronuclear stages in zygotes treated with Flavopiridol. (F) Quantification of the percentage of zygotes treated with Flavopiridol with maternal CENP-A-mScarlet deposited in both pronuclei or only in the maternal pronuclei at each pronuclear stage. G) Quantification of pronuclear stages in zygotes treated with BI2536. H) Quantification of the percentage of zygotes with maternal CENP-A-mScarlet in both pronuclei or only in the maternal pronuclei after 8hrs of fertilization in control, Flavo-treated, Flavo-treated PN3-only, and BI2536-treated embryos.

