## Supplemental Figures #1-7 for "Equalizing epigenetically imprinted centromeres in early mammalian embryos"

### Supplemental Figure 1:

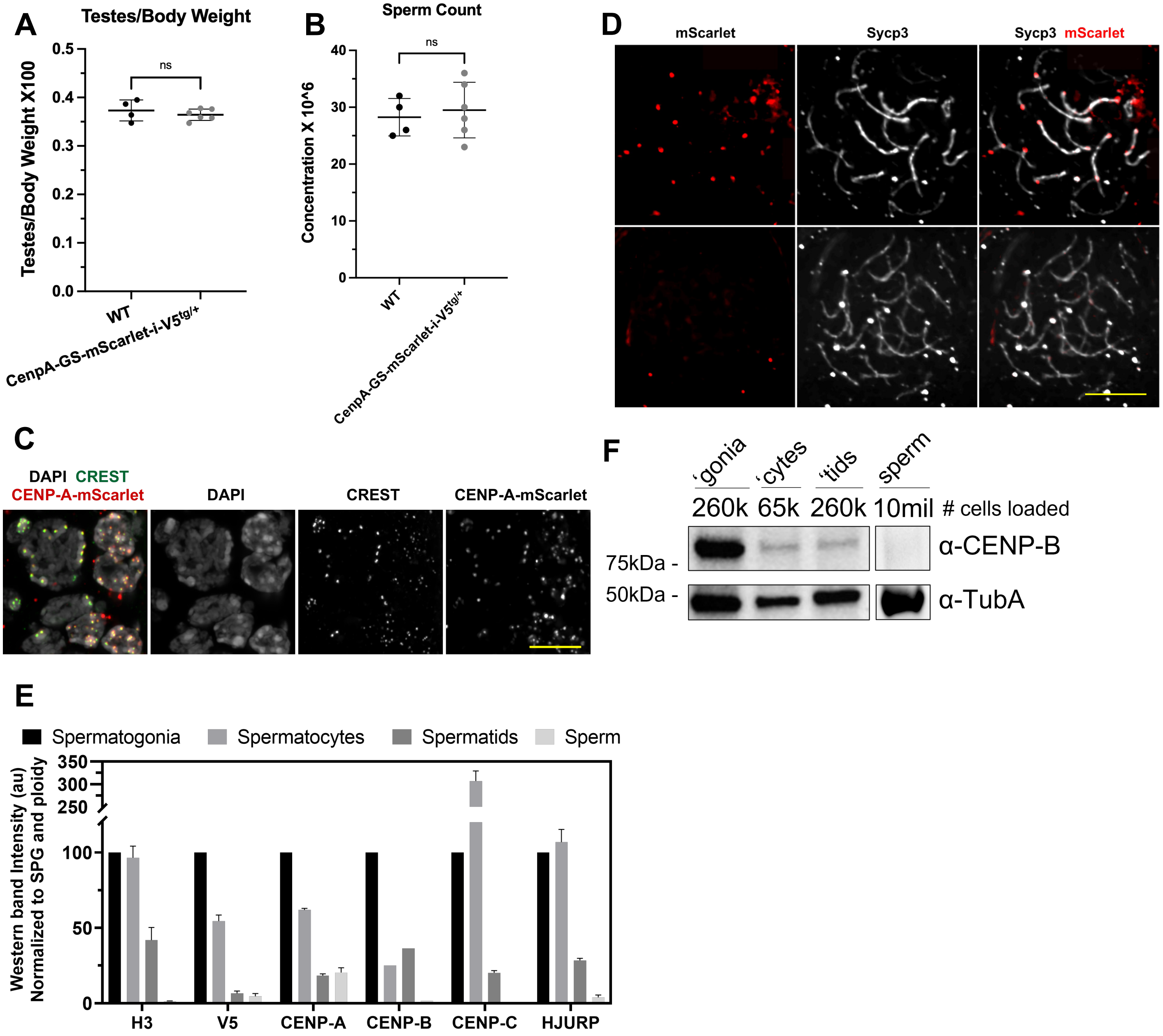

#### Supplemental Figure 2:

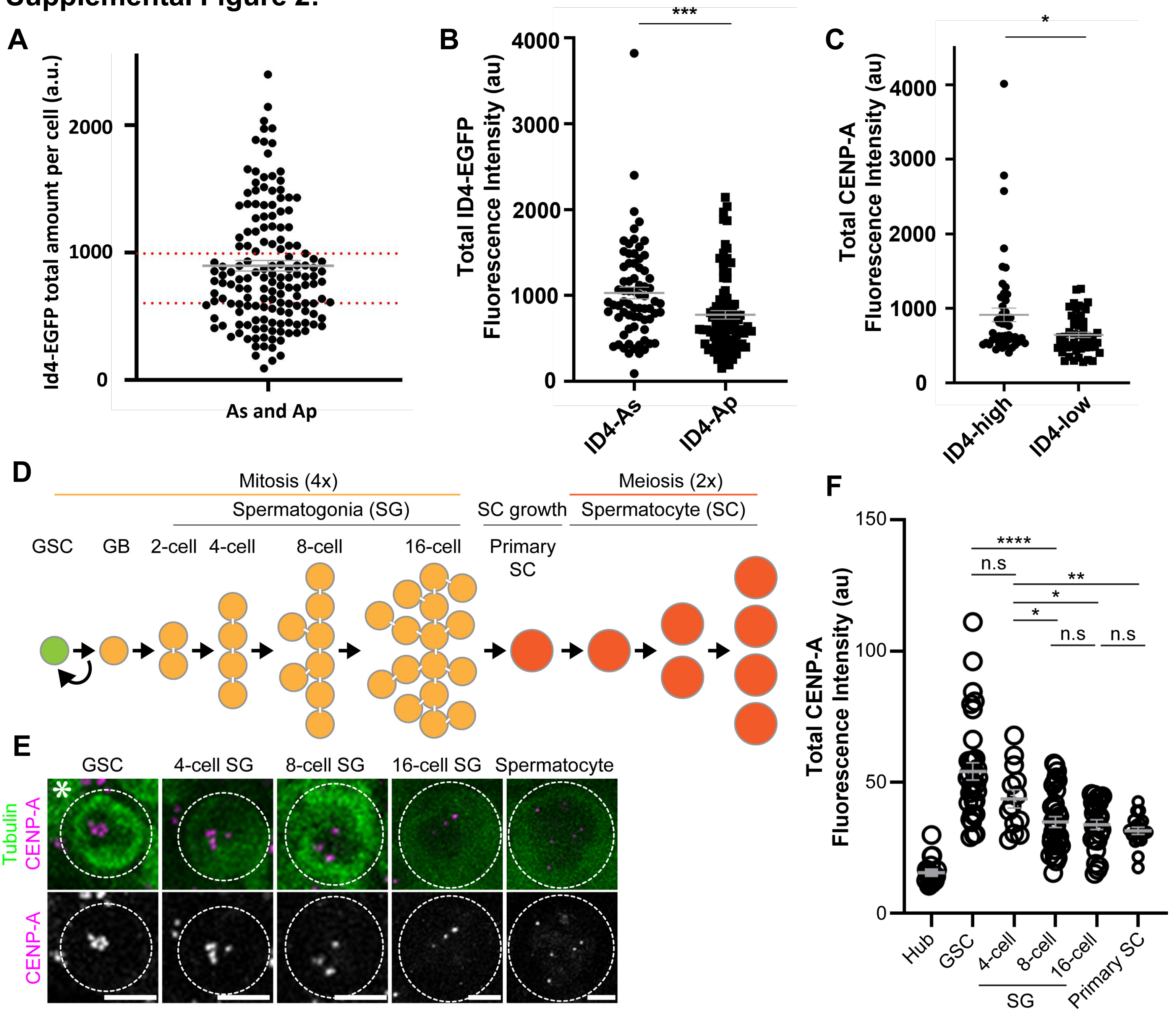

#### Supplemental Figure 3:

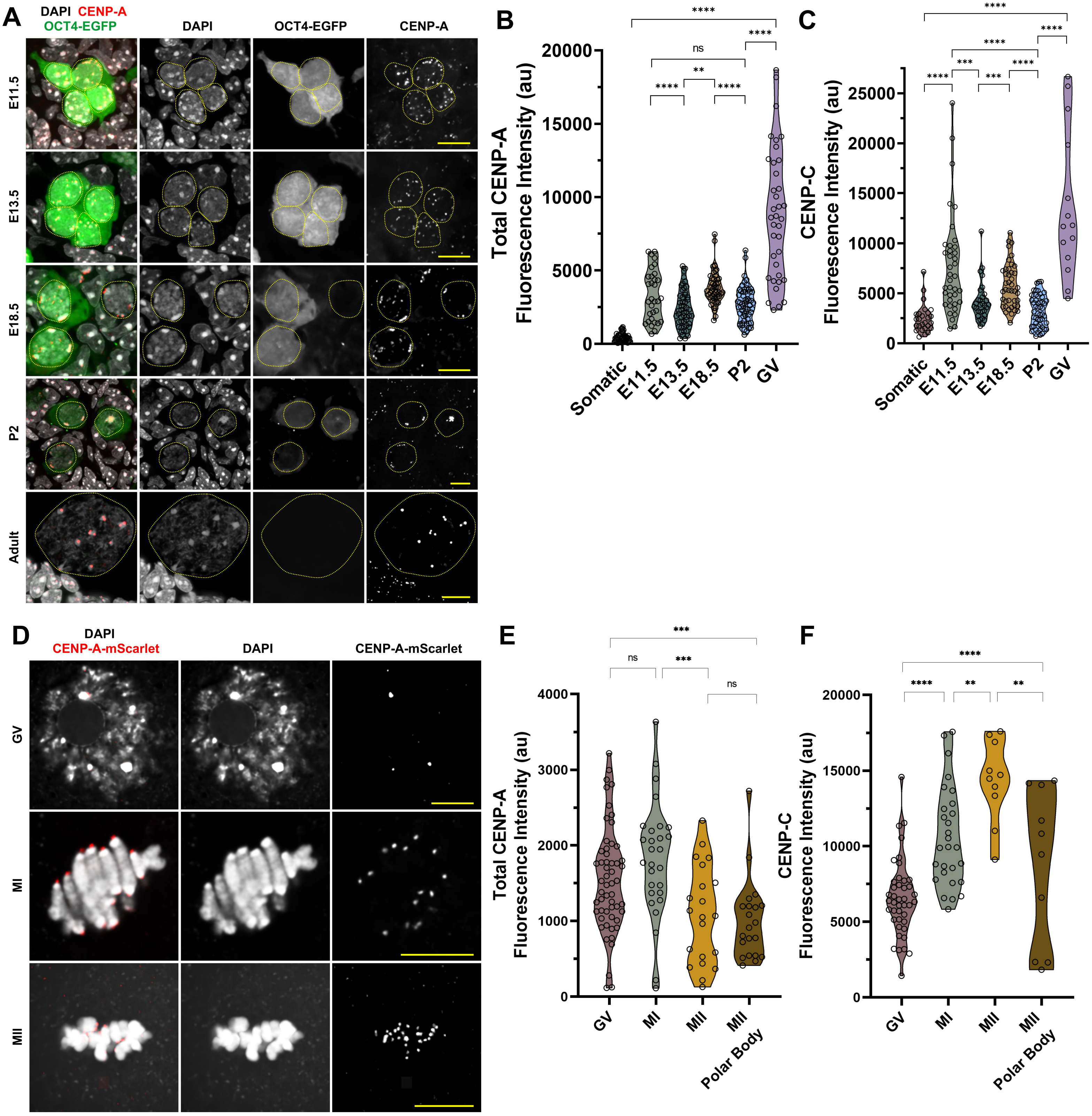

#### Supplemental Figure 4:

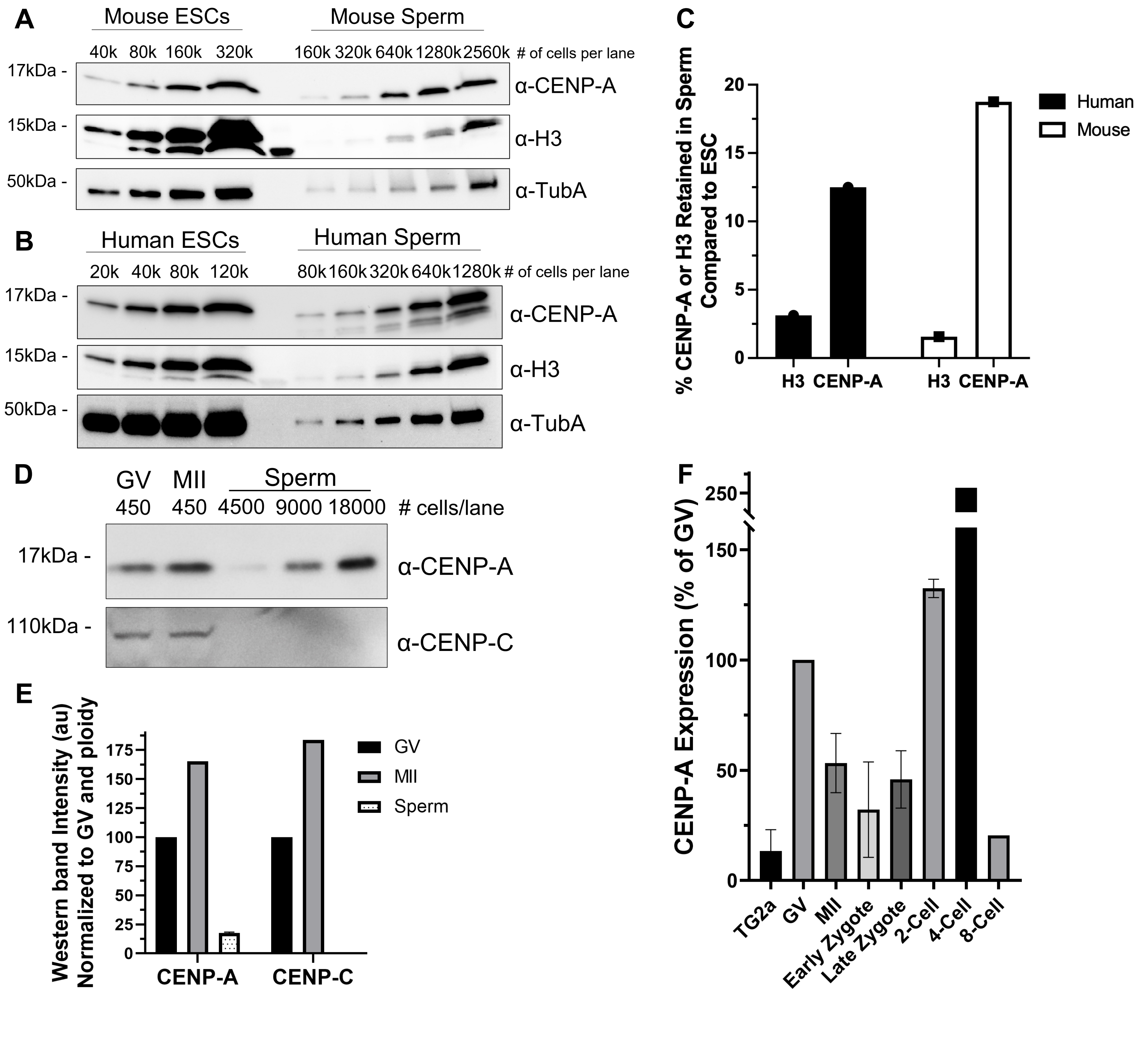

### Supplemental Figure 5:

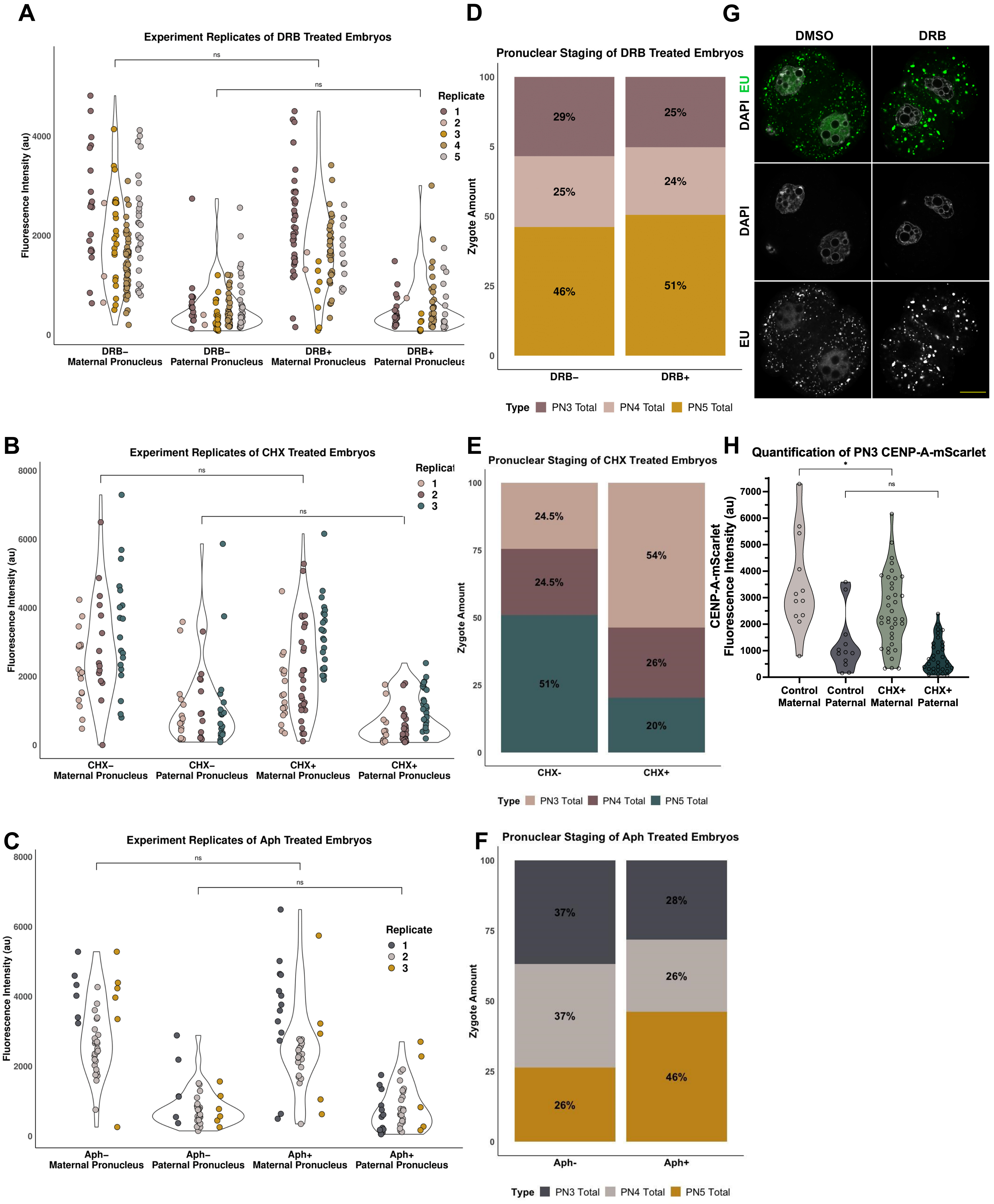

#### Supplemental Figure 6:

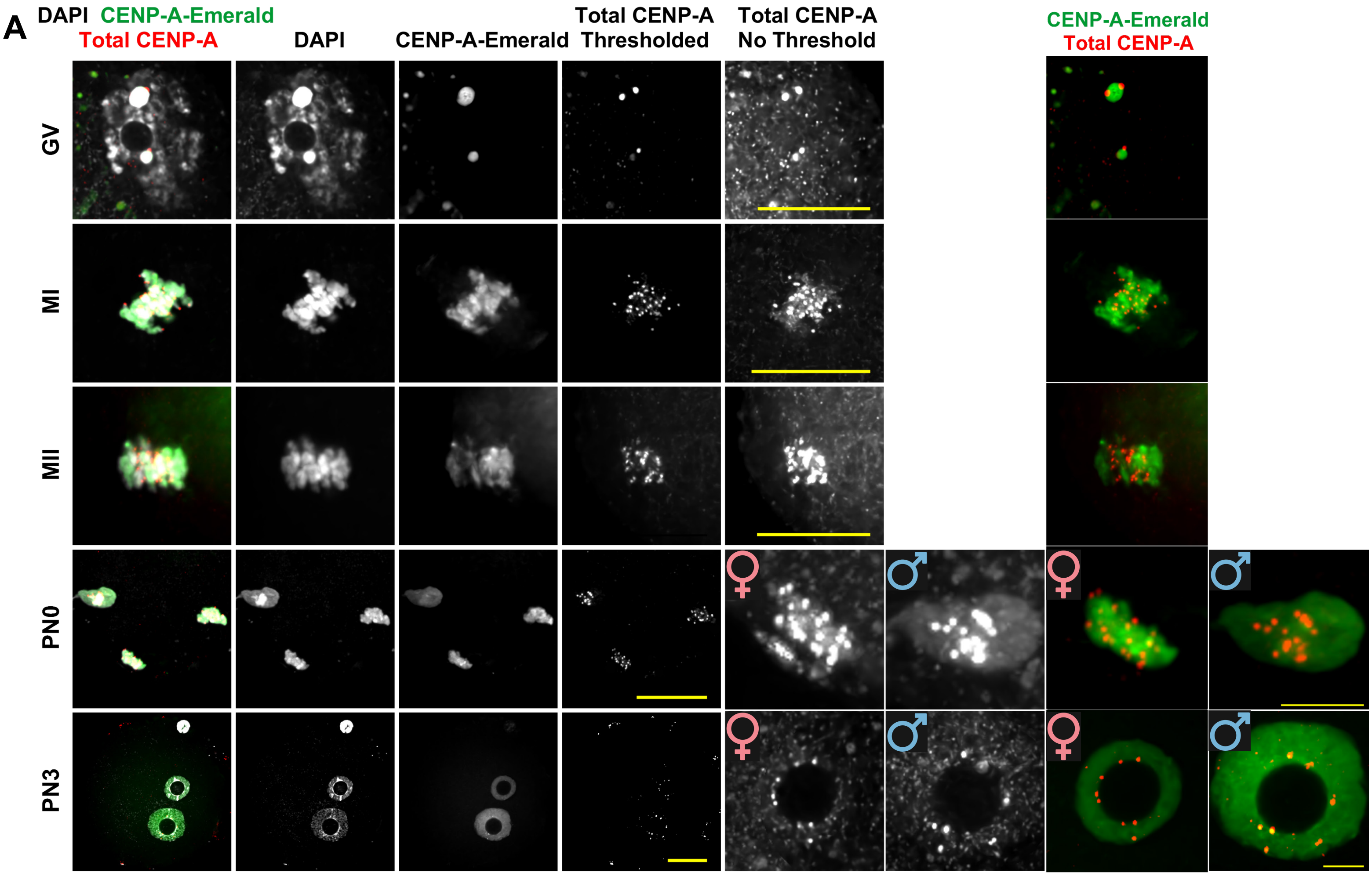


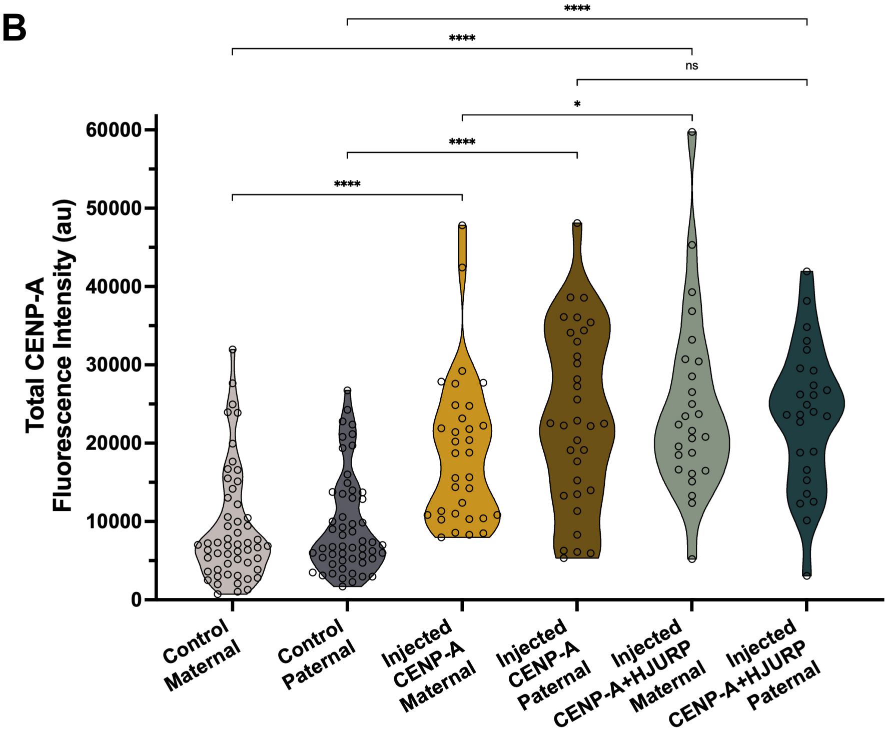

#### Supplemental Figure 7:

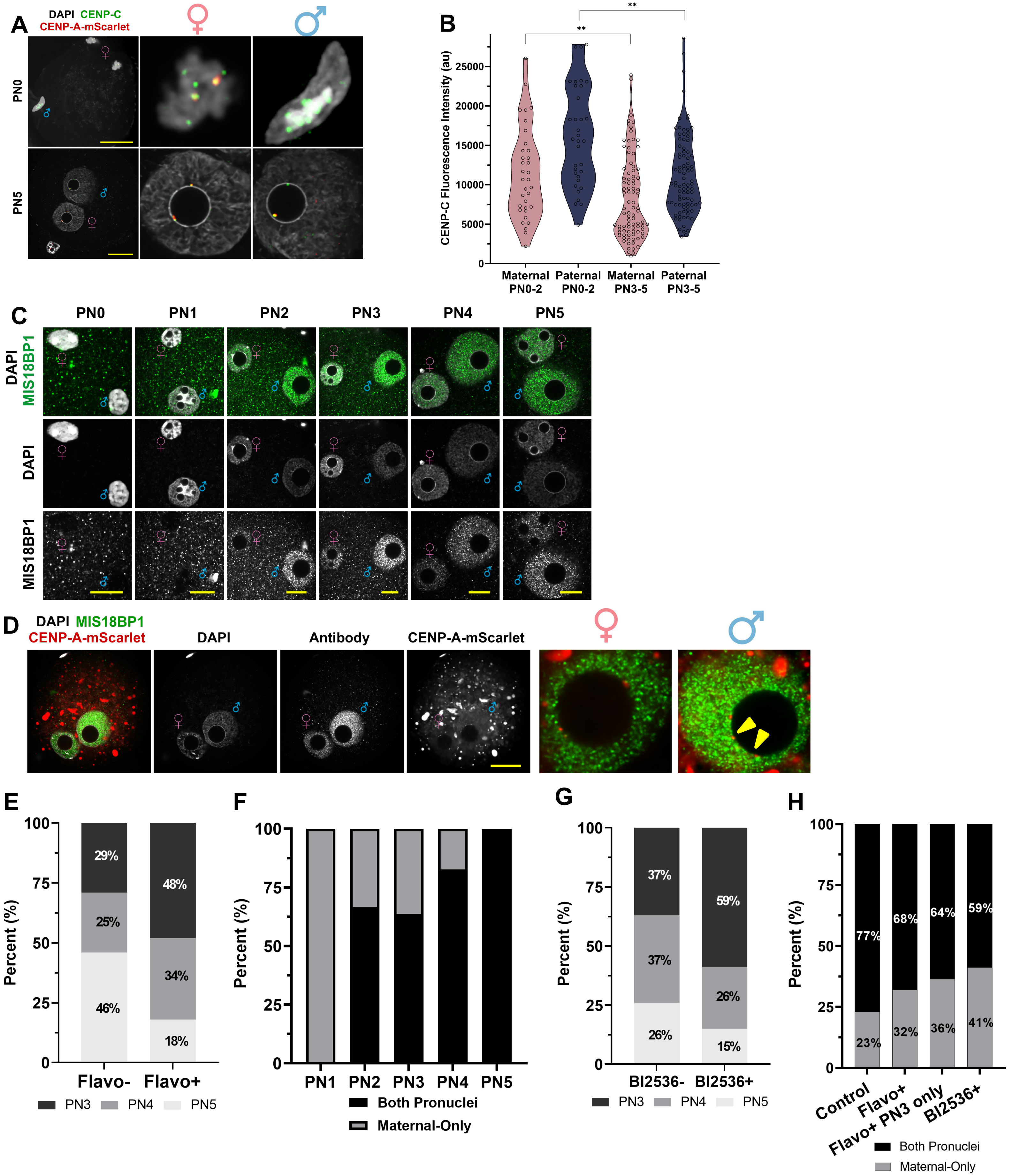
